# AI-assisted improvement of *Aspergillus oryzae* β-galactosidase using an Ensemble of Protein Language Models

**DOI:** 10.64898/2026.05.20.726739

**Authors:** Andrea Trapote Fernández, Alicia Fernández, Juan A. Méndez-Líter, Alicia Prieto, Jorge Barriuso, Fernando G. Osorio

## Abstract

β-galactosidases (BGs) are essential enzymes widely used in the food industry, particularly in the production of lactose-free products. Among them, the BG from *Aspergillus oryzae* is of industrial relevance due to its activity at acidic pH and moderate thermal tolerance. However, enhancing its catalytic performance remains a key challenge. Traditional enzyme engineering methods are time-consuming and resource-intensive, limiting their scalability. Recent advances in Artificial Intelligence (AI), particularly those based on Natural Language Processing, offer a promising alternative by enabling efficient exploration of protein sequence space and prediction of beneficial mutations. In this study, we introduce an ensemble-based, zero-shot Protein Language Model pipeline that reconciles predictions from six independent models (ESM2 and the five ESM1v variants) combined with a diversity-aware candidate selection strategy. Applied to the BG from *A. oryzae*, this approach identified beneficial mutations leading to novel enzyme variants with up to a four-fold increase in catalytic efficiency on oNPGal, a two-fold increase on lactose, and, independently, a T338I variant with markedly enhanced thermostability (≈80% residual activity after 24 h at 60 °C), all without requiring supervised fine-tuning on experimental fitness data. Our results demonstrate that consensus across an ensemble of PLMs can efficiently enrich beneficial substitutions in industrially relevant enzymes and substantially reduce the number of wet-lab candidates that need to be screened.

**Table of Contents graphic:** 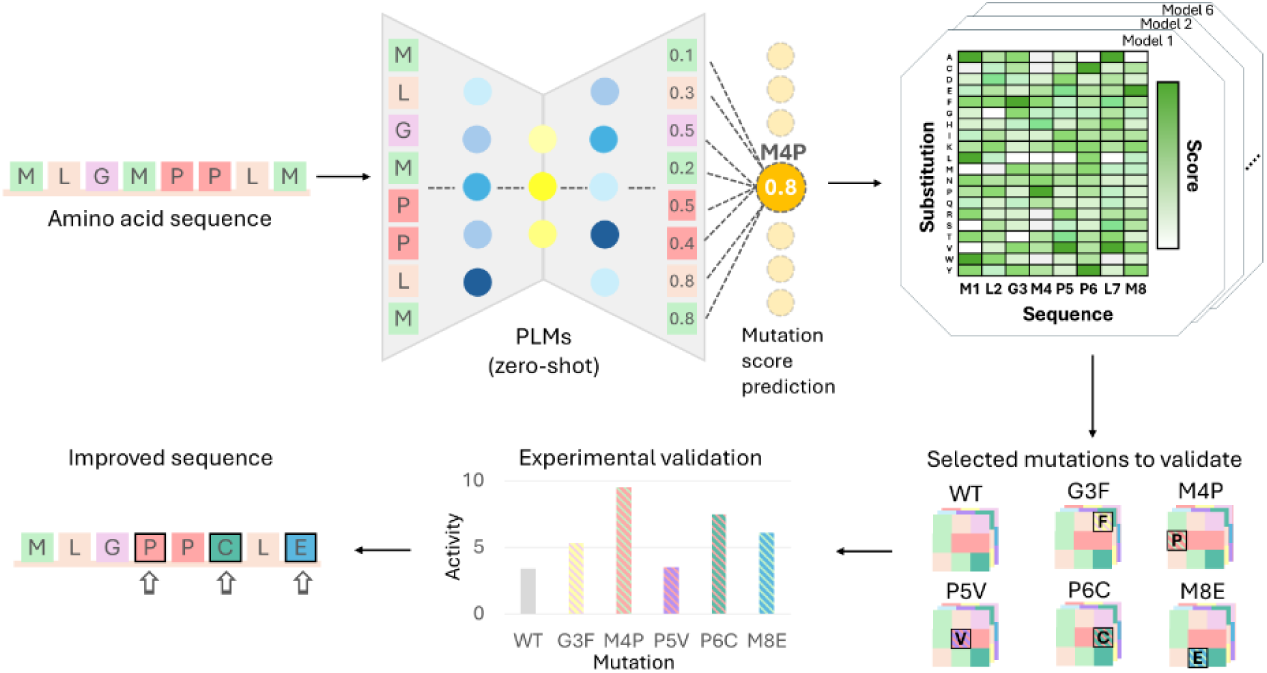

## INTRODUCTION

Enzymes are indispensable catalysts in modern industry, facilitating a wide array of chemical reactions with remarkable efficiency and minimal environmental impact^1^. These biocatalysts find applications across various sectors, including food, detergents, biofuels, textiles, and pharmaceutical production^2^. Within the food industry, enzymes play a pivotal role in enhancing food safety and developing innovative products. For instance, amylases improve bread texture and flavor^3^, while proteases are used in cheese manufacturing to accelerate ripening and develop flavor^4^.

Among these important industrial enzymes, β-galactosidases (EC 3.2.1.23), commonly known as lactases^5^, are glycoside hydrolases (Carbohydrate-Active enZymes, CAZymes) particularly important due to their role in hydrolyzing lactose into glucose and galactose. Produced by various microorganisms, their physicochemical properties and substrate specificity vary depending on the source. Most of these enzymes are classified into different glycosyl hydrolase (GH) families, including: GH1, which usually groups thermophilic lactases from bacteria and archaea; GH2, mainly from bacteria and yeasts; GH35, primarily fungal; and GH42, usually bacterial cold-adapted enzymes^6^.

The enzymatic reaction catalyzed by BGs is crucial for producing lactose-free dairy products for individuals with lactose intolerances^7–9^. Traditionally, industrial enzymes have been sourced from nature and subsequently engineered to enhance their efficiency and withstand industrial process conditions^10^. However, classical enzyme engineering, involving mutagenesis and screening, is a labor-intensive and time-consuming process that often requires testing thousands of variants, thus limiting progress^11,12^. Even before the emergence of modern artificial intelligence (AI)-based methods, computational design approaches had already demonstrated their ability to improve protein stability and expression through structure and sequence guided optimization^13^. Recent advancements in AI have revolutionized enzyme engineering^14^. Machine learning (ML) and deep learning (DL) techniques can analyze vast datasets to identify patterns and predict the effects of mutations on enzyme properties^15–19^. One notable application is AlphaFold, which enables accurate protein folding predictions, facilitating the understanding of structure-function relationships^20,21^. In this regard, generative AI models, particularly those employing Natural Language Processing (NLP) techniques, can design novel enzyme sequences with enhanced or entirely new functionalities^14^, offering a more efficient and rapid approach to classical rational design and directed evolution approaches through the prediction and selection of beneficial mutations.

Very recently, a type of NLP technique known as Protein Language Models (PLMs) has emerged as a powerful paradigm for protein sequence optimization^22–24^. Inspired by their success with textual data, PLMs treat amino acids as a twenty-element vocabulary, enabling them to capture intricate patterns and contextual dependencies within protein sequences, thus facilitating efficient exploration and optimization of sequence space.

Our work builds on the established idea that PLMs, trained on massive sequence datasets, capture evolutionary constraints that correlate with protein fitness. Based on this principle, we postulate that high-score mutations identified by integrated PLMs are more likely to produce structurally stable variants, thereby increasing the chances of observing improved catalytic properties compared to random mutagenesis. In the context of β-galactosidases, AI-based approaches have been scarcely employed to enhance their thermal stability and catalytic efficiency^25^. By training models on existing enzyme datasets, researchers can predict mutations that confer resistance to higher temperatures, a desirable trait for industrial applications.

Recent PLM-guided protein engineering efforts have pursued two broadly distinct strategies: supervised pipelines that couple PLM-derived representations with machine-learning regressors fitted on experimental fitness data¹⁷,¹⁸,⁴², and zero-shot consensus approaches, pioneered in antibody engineering⁴³, that rank substitutions by evolutionary likelihood without retraining. Enzyme engineering raises challenges that are not present in antibody optimization: (i) the catalytic machinery is concentrated in a small, highly conserved region whose residues must be preserved, whereas antibody diversification deliberately targets the complementarity-determining regions; (ii) the optimization endpoint is a kinetic parameter, not a binding affinity, and evolutionary likelihood does not map monotonically onto catalytic rate; (iii) industrial enzymes must simultaneously satisfy multiple physicochemical objectives — activity, thermostability, pH tolerance, substrate specificity. Here we extend the zero-shot ensemble logic to enzyme engineering through a conciliation scheme across six independent PLMs and a diversity-aware selection step that balances exploitation of high-consensus substitutions with exploration across sequence domains.

In this study, we focus on the optimization of β-galactosidase produced by the fungus *Aspergillus oryzae* (GH35), a crucial enzyme for producing lactose-free foods^27^. β-Galactosidase from *A. oryzae* is highly suitable for industrial applications due to its optimal activity at acidic pH (around 4.5 to 5) and temperature tolerance up to 50°C, aligning with common food processing conditions^28^. Additionally, the β-galactosidase from *A. oryzae* is recognized for its significant transgalactosylation activity, resulting in the production of galactooligosaccharides (GOS), valuable prebiotic compounds^29–32^.

By leveraging PLM models to predict and design BGs with enhanced catalytic efficiency, we accelerate the engineering of β-galactosidase enzymes. In this work, we implement a computational framework based on PLMs that enables adaptation of enzymes to diverse industrial conditions, ultimately yielding a novel β-galactosidase with improved catalytic efficiency. This approach can be applied not only to produce BGs with superior properties, but also to establish a methodological framework for sequences optimization applicable to industrial enzymes.

## MATERIALS AND METHODS

### Selection of the model enzyme

A screening of BG sequences of industrial relevance was conducted by searching for sequences of interest across various public sources, including GenBank and UniProt databases^33,34^, research articles, reviews, and patents. Only biochemically characterized enzymes were considered, to avoid possible bias of sequencing annotations, and to be sure of the different properties of the selected enzymes. A total of 45 candidate sequences were evaluated based on their functional properties, including substrate specificity, thermal stability, and quaternary structure, spanning lactases from different families (GH1, GH2, GH35, and GH42).

### Protein language model-based prediction

*A. oryzae* β-galactosidase was optimized using state-of-the-art Protein Language Models, specifically ESM2 and ESM1v^35,36^. ESM2, an advanced iteration of the original ESM model, uses deeper network architecture and extensive pretraining on large evolutionary sequence databases, which positions this PLM as a standard in the field. In contrast, ESM1v comprises an ensemble of five models specifically fine-tuned for variant effect prediction. By aggregating predictions from these multiple models, ESM1v enhances the robustness and accuracy of its predictions, mitigating bias and variability, and offering a more holistic interpretation of protein sequences.

Mutation effects were estimated using the *masked marginal scoring* procedure implemented in these PLMs. Given a wild-type (WT) sequence *x*, each mutated position *i* is replaced by a special *<mask>* token while keeping the rest of the sequence unchanged. The masked sequence is then processed by the model, which outputs a vector of logits at each position. These logits are transformed into a probability distribution (*P*) over amino acids at each masked position conditioned on the sequence context using the softmax function. For each mutation at position *i*, the effect of substituting the wild-type residue *wt*_*i*_ with a mutant residue *mt*_*i*_ is quantified as the difference in log-probabilities:

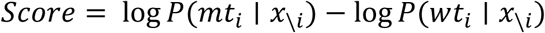

This score represents the predicted change in sequence log-probability under the model due to the specified mutations. Positive values indicate that the mutant residues are more compatible with the sequence context than the wild-type residues according to the model, while negative values indicate the opposite.

To improve robustness, we used an ensemble of six pretrained models (ESM2 and ESM1v_1-5)._ Initially, we ran each PLM to estimate position-specific substitution score for every amino acid at each position in the initial sequence. We then selected a candidate substitution for each position from these PLM-derived score using a combination criterion. This criterion selected mutations consistently favored across all models and ranked them based on the normalized average of their outputs (Figure 1). This integration step yielded a candidate group of unique substitutions for each position.

**Figure 1.**
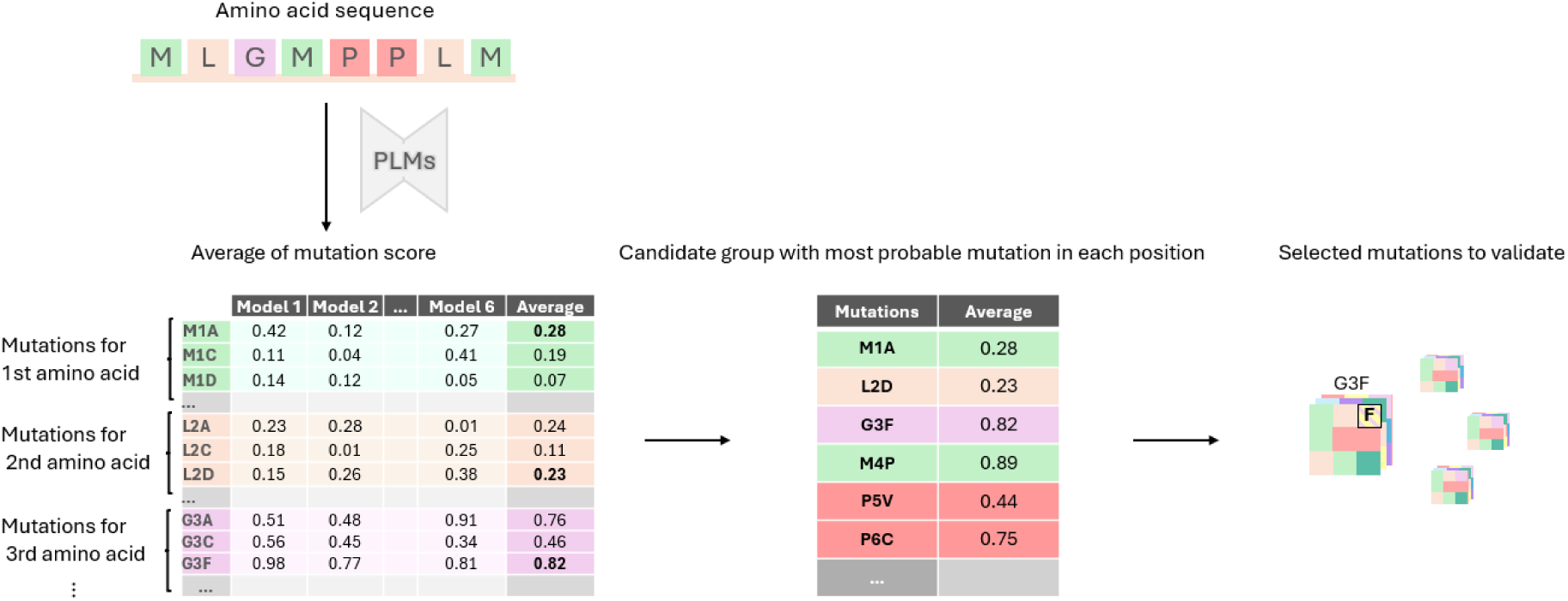
Workflow of the six-model PLM ensemble pipeline. Each position of the wild-type sequence is masked and processed independently by ESM2 and the five ESM1v models. For each model, the masked-language-modelling head produces a 20-dimensional score over amino acids. Per-position scores are averaged across the six models and mutations are ranked by the resulting consensus score. Candidates for experimental validation are then selected combining the top consensus leaders with additional high-scoring mutations from under-represented sequence domains (diversity-aware step).

Then, we implemented a two-step systematic strategy to identify seven candidates for experimental validation. First, we selected the top four mutations with the highest average score across all six PLMs (consensus leaders). Second, to promote diversity, three additional high-scoring mutations were chosen from sequence regions not represented in the initial set. This approach balanced the exploitation of high-confidence predictions with the exploration of diverse sequence domains.

### Prediction of the 3D structures of β-galactosidases

The amino acid sequence from the wild-type and the 7 mutants was modeled using AlphaFold2^20,21^ and the resulting structure was visualized using 3Dmol to assess the position of each of the seven proposed mutations. Additionally, Caver Analyst 2.0 was used to model the catalytic pocket of the enzymes^37^, along with the amino acids that form it.

### Heterologous expression of β-galactosidase variants

The 7 mutant genes of the selected BGs, derived from the fungus *A. oryzae* sequence, were codon optimized for expression in the heterologous host and synthesized by GeneScript (Table S1). The wild-type version, with the original sequence of the native protein, and seven different mutants were produced and directly cloned into pPIC9 plasmid using EcoRI and NotI restriction enzymes (Invitrogen), suitable for expression in the yeast *Komagataella phaffii* (syn. *Pichia pastoris*). All recombinant constructs included the *Saccharomyces cerevisiae* α-factor signal peptide for secretion in *K. pastoris*. Heterologous expression of the genes for the eight β-galactosidases was carried out using the *K. pastoris* GS115 strain (Invitrogen), which was maintained on YPD agar plates (10 g/L yeast extract, 20 g/L peptone, 20 g/L glucose, and 10 g/L agar). Yeast transformation was performed by electroporation, following the protocol outlined in the EasySelect™ *Pichia* Expression Kit (Invitrogen, Waltham, MA, USA), introducing a plasmid containing the gene for the corresponding β-galactosidase and the HIS selection marker. Transformants were selected on YNΒ-His agar plates (20 g/L glucose, 6.7 g/L YNB, 1.92 g/L histidine-free amino acids (Sigma-Aldrich), and 10g/L agar), and were cultured for up to 72h at 28°C. Finally, for recombinant protein production, YEPS medium (20 g/L peptone, 10 g/L yeast extract, 10 g/L sorbitol, and 100mM potassium phosphate buffer, pH 6) was used, with daily addition of 1% methanol as an inducer. The cultures were incubated for 4 days at 28 °C and 250 rpm, with samples taken daily to measure protein production.

### Enzyme production and purification

Purification of the eight β-galactosidase variants was carried out using an ÄKTA Purifier FPLC system (GE Healthcare Life Sciences, Chicago, IL, USA). First, the yeast cells were removed by centrifugation, and the supernatant was clarified using sequential filtration processes, ending with a 0.2µm filter.

Subsequently, the enzymes were dialyzed against 10 mM phosphate buffer, pH 6, containing 100mM NaCl, and were purified in a single chromatographic step by size-exclusion chromatography using a HiLoad 16/60 Superdex 200 column (GE Healthcare Life Sciences)^28^. To prevent nonspecific interactions, 100mM NaCl was used to equilibrate the column and elute proteins at a flow rate of 1mL/min.

Fractions containing β-galactosidase activity were collected, concentrated, dialyzed to remove NaCl, and their purity was confirmed by SDS-PAGE. The theoretical molecular weight of the mature β-galactosidase polypeptide (excluding the signal peptide) is approximately 108kDa, and the molar extinction coefficient at 280nm is ε₂₈₀ = 1.92 × 10⁵M⁻¹cm⁻¹, as calculated from the amino acid sequence.

### Sedimentation velocity assay (SV)

Samples of WT enzyme and E460G (dissolved in sodium acetate buffer, 10 mM pH 5) were loaded (320 µL) into analytical ultracentrifugation cells. The experiments were carried out at 20 °C and 48000 rpm in an XL-A analytical ultracentrifuge (Beckman-Coulter Inc.) equipped with UV-VIS absorbance detection systems, using an An-50Ti rotor, and 12 mm Epon-charcoal standard double-sector centerpieces. Sedimentation profiles were recorded at 280 nm. Differential sedimentation coefficient distributions were calculated by least-squares boundary modelling of sedimentation velocity data using the continuous distribution c(s) Lamm equation model as implemented by SEDFIT^38^. Experimental values were corrected to standard conditions (water, 20 °C, and infinite dilution) using the program SEDNTERP^39^ to get the corresponding standard s values (s20,w).

### Protein quantification, activity assays and substrate specificity

The protein concentration of the purified enzymes was determined by measuring absorbance at 280 nm using a Nanodrop spectrophotometer. This value was subsequently used to calculate their specific activity.

The standard assay for β-galactosidase activity was carried out using 0.1% (w/v) *o*-nitrophenyl-β-D-galactopyranoside (*o*NPGal, Sigma) as the substrate, a lactose analogue, at 50 °C in 100 mM sodium acetate buffer. For *A. oryzae* β-galactosidase, reactions were buffered at pH 5 with 50 mM sodium acetate buffer, as previously described^40^. Reactions were stopped after 10 min by adding 2% (w/v) Na_2_CO_3_, and the released *o*NP was measured spectrophotometrically at 410 nm. One unit of BG activity was defined as the amount of enzyme capable of releasing 1 micromole of *o*NP per minute (the molar extinction coefficient of *o*NP is 15,200 M^-1^cm^-1^).

Once the activity of the β-galactosidase from *A. oryzae* was studied over *o*NPGal, the hydrolysis of lactose was evaluated. The activity of β-galactosidase on lactose was quantified by measuring the glucose released from these compounds after enzymatic hydrolysis, using the commercial Glucose-TR kit (Spinreact), following the manufacturer’s instructions. Because lactose hydrolysis yields equimolar amounts of glucose and galactose, the measured glucose concentration directly corresponds to the amount of lactose hydrolyzed and was used without further correction. The reactions were carried out in 100 mM sodium acetate buffer, pH 5, incubated in a thermal block for 10 min at 1200 rpm. The reactions were then stopped by inactivating the enzyme at 100 °C for 10 min.

The kinetic constants of the purified enzymes were determined using *o*NPGal (ranging from 0.2 mM to 60 mM) and lactose (ranging from 1.4 mM to 250 mM). The *K_m,_ V_max_* and *k_cat_* values, and catalytic efficiency of the enzymes (*k_cat_/K_m_*) were determined using SigmaPlot 15.0 software, based on the Michaelis-Menten model. All experiments were performed at the optimal pH and temperature of the enzymes. Prior to the determination of kinetic parameters, reaction linearity was confirmed by measuring product formation at 5, 10, and 20 min under the standard assay conditions described above for all enzyme variants. The corresponding activity values (U/mL) for each time point are provided in Table S2. All Michaelis–Menten curves used to derive kinetic parameters have been included in the Supporting Information (Figures S7-S8), together with the corresponding nonlinear regression fits, R² values, and standard deviations.

### Physicochemical properties of the enzymes

The effect of pH on enzymatic activity was evaluated using 100 mM Britton–Robinson buffer, which allows assay conditions to be adjusted over a broad pH range (2–10). Optimal pH values were determined by incubating purified WT and mutant β-galactosidases (H83Q, S137P, Y316A, T338I, E460G, S783P, and Q867L) in this buffer at a specific pH with *o*NPGal as substrate for 10 min, at 1200 rpm, and 50 °C. To assess pH stability, enzyme samples were incubated in Britton–Robinson buffer adjusted to pH values from 2 to 10 for 24 h at 4°C. Residual activity was then measured under standard β-galactosidase assay, at pH 5.

Thermostability was examined by incubating purified enzymes for 24 h at 30, 40, 50, 60 and 70 °C in 100 mM sodium acetate, pH 5. After incubation, residual activity was measured under standard assay conditions. All treatments were performed in triplicate.

### HPLC analysis of transgalactosylation products

The lactose hydrolysis products were also analyzed by high-performance liquid chromatography (HPLC), using an Agilent 1200 series LC instrument. The Aminex HPX-42C column was employed, under the operating conditions described by the manufacturer, with MilliQ H_2_O as the mobile phase. The product peaks were detected by their refractive index (RI) and identified by comparing their retention times with those of lactose, glucose, and galactose standards analyzed under identical conditions.

Transgalactosylation reactions were performed using 9.4 µg of each enzyme in 20 mM acetate buffer (pH 5). The final reaction volume was 400 µL. A 250 mM concentration of lactose was used, as high substrate concentrations favor transgalactosylation reactions over hydrolysis processes. Transgalactosylation products were analyzed using the same procedure explained for lactose hydrolysis analysis. Reactions were maintained for 5h, with samples taken at 20 min, 1 h, 2 h, and 5h. Quantification was expressed in lactose equivalents, as commercial standards for the different galacto-oligosaccharides were not available. Although commercial standards for these compounds are not available, the observed chromatographic behavior is consistent with the retention patterns described in the literature for β-(1→3) and β-(1→6) linked GOS.

## RESULTS AND DISCUSSION

### Enzyme selection

As a first step in the definition of an AI-based framework for optimization of enzymes, a systematic survey of β-galactosidase sequences of industrial relevance was conducted. Sequences were evaluated based on their functional properties, including factors such as their secondary, tertiary and quaternary structure, substrate specificity, thermal stability, and the potential for effective lactose hydrolysis. Derived from this analysis, the *A. oryzae* β-galactosidase (UniProt: Q2UCU3) was selected due to its promising characteristics in terms of activity, robustness, and monomeric architecture.

Monomeric enzymes, with their single-subunit architecture, offer a simplified system for analysis and modelling, facilitating a clearer understanding of structure-function relationships and the impact of mutations. Their reduced complexity also streamlines experimental analysis, such as evaluating the effects of specific mutations on activity or stability^41^. The monomeric architecture of A. oryzae β-galactosidase additionally simplifies the attribution of phenotypic effects to individual residues, since confounding inter-subunit allosteric effects are absent.

### PLM-based prediction

The use of PLMs for enzyme optimization has been previously reported^14,26,42^, yet their predictive accuracy in this domain has generally been lower compared to applications such as PLM-driven antibody optimization^43,44^. Previous strategies often required substantial experimental validation, sometimes incorporating ML regressors to correlate PLM predictions with functional data^18^. To minimize the reliance on extensive experimental screening and adhere to a zero-shot inference framework—where models are not fine-tuned on specific experimental fitness landscapes—we designed a prediction workflow that integrates the outputs of multiple PLMs and employs *in silico* conciliation of their predictions.

In this study, the wild-type sequence of *A. oryzae* β-galactosidase was analyzed using six distinct PLM models: ESM2 and five variants of ESM1v. For each model, we obtained the full 20-dimensional score distribution over amino acids at each position of the sequence via the masked-language-modelling head, as detailed in Materials and Methods. Predictions from the six models were combined by averaging the normalized amino acid substitution score, yielding a consensus score for each candidate substitution across the ensemble (Figure S1 and Table S3). The underlying hypothesis behind this consensus strategy is that substitutions consistently favored across an ensemble of independently trained PLMs are more likely to preserve structural integrity and may enhance catalytic performance, while individual model biases are averaged out.

The implementation of this AI approach presents a notable advantage over traditional protein design methodologies. It facilitates a more detailed analysis and exploration of the intricate relationships between sequence, structure, and function, thereby enabling the design of enzyme variants with optimized properties^45^. In contrast, this AI-driven strategy explores a wider sequence space, enabling the design of novel variants that are difficult to access through traditional experimental approaches.

### Experimental validation of PLM-predicted substitutions

We selected seven substitutions for subsequent experimental validation based on their average substitution score (Table 1 and Figure 2), ensuring coverage across the entire primary sequence and with the expectation of obtaining improved properties of the enzyme. Table 1 shows the scores assigned by the six PLMs to each selected mutation, as well as their average values, while Table 2 presents the ranking positions of each selected mutation for each PLM according to its score, together with their average ranking.

**Figure 2.**
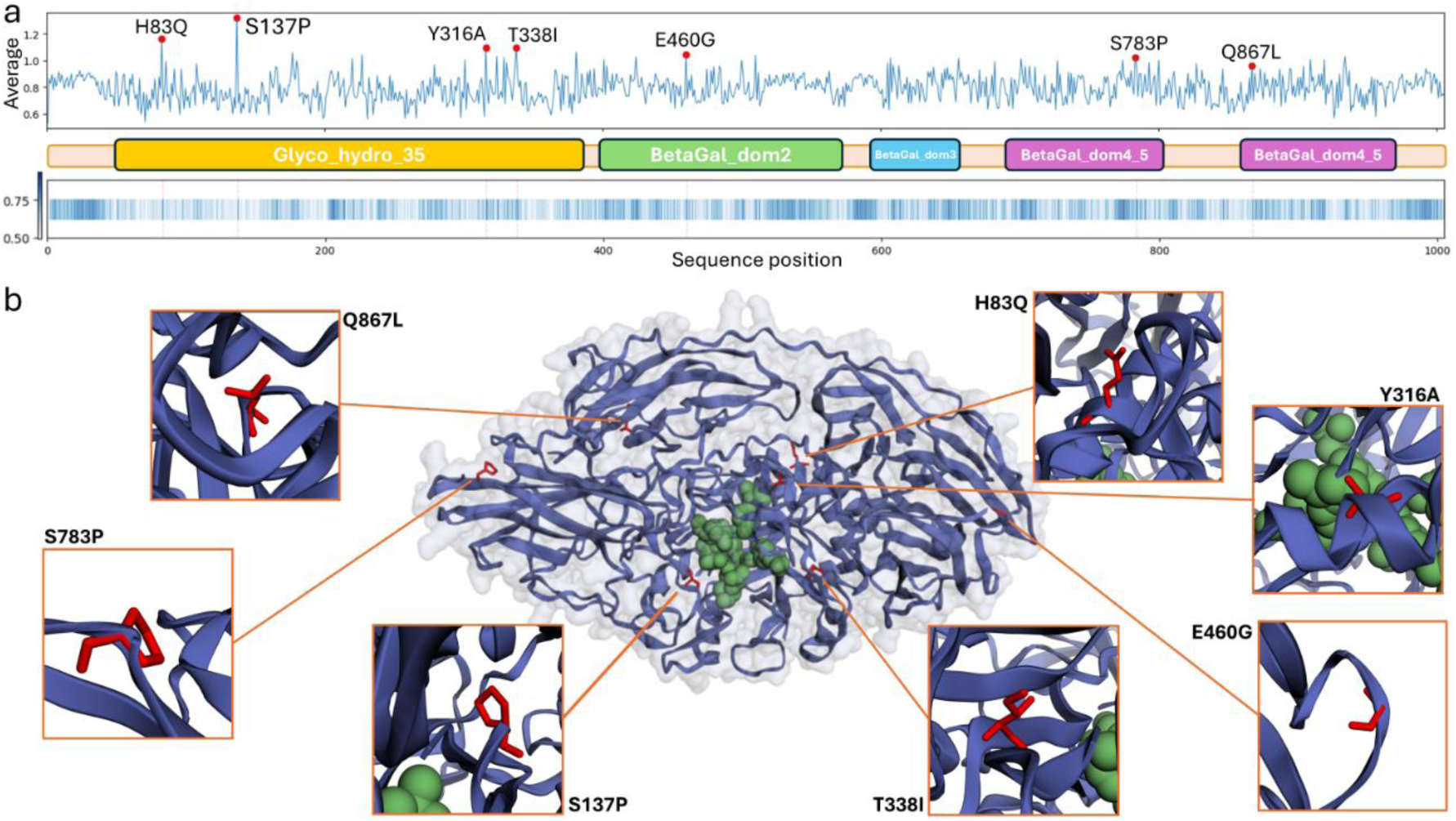
(a) The sequence of A. oryzae β-galactosidase is represented with its specific domains highlighted in different colors, along with the average normalized substitution score at each position. The seven selected mutations are marked in red. (b) Mapping of the mutations in the AlphaFold2 predicted structure. The catalytic pocket of the enzyme is highlighted in green and the localization of the selected amino acid substitutions: H83Q, S137P, Y316A, T338I, E460G, S783P, and Q867L in red.

**Table 1.**
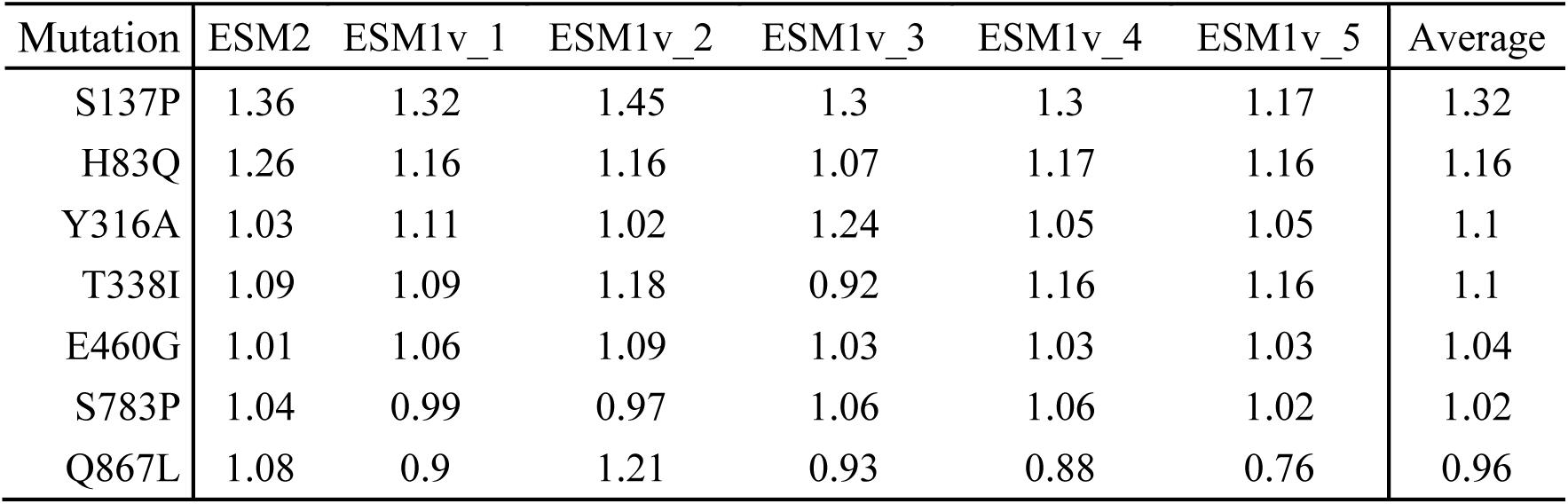
List of *A. oryzae* β-galactosidase selected substitutions and predicted scores across the different PLMs and their average.

| Mutation | ESM2 | ESM1v_1 | ESM1v_2 | ESM1v_3 | ESM1v_4 | ESM1v_5 | Average |
| --- | --- | --- | --- | --- | --- | --- | --- |
| S137P | 1.36 | 1.32 | 1.45 | 1.3 | 1.3 | 1.17 | 1.32 |
| H83Q | 1.26 | 1.16 | 1.16 | 1.07 | 1.17 | 1.16 | 1.16 |
| Y316A | 1.03 | 1.11 | 1.02 | 1.24 | 1.05 | 1.05 | 1.1 |
| T338I | 1.09 | 1.09 | 1.18 | 0.92 | 1.16 | 1.16 | 1.1 |
| E460G | 1.01 | 1.06 | 1.09 | 1.03 | 1.03 | 1.03 | 1.04 |
| S783P | 1.04 | 0.99 | 0.97 | 1.06 | 1.06 | 1.02 | 1.02 |
| Q867L | 1.08 | 0.9 | 1.21 | 0.93 | 0.88 | 0.76 | 0.96 |

**Table 2.** Ranking of the selected mutations across the different PLMs. 1 indicates the highest score. The average column represents the re-ranking based on the consensus score.

| Mutation | ESM2 | ESM1v_1 | ESM1v_2 | ESM1v_3 | ESM1v_4 | ESM1v_5 | Average |
| --- | --- | --- | --- | --- | --- | --- | --- |
| S137P | 1 | 1 | 1 | 1 | 1 | 1 | 1 |
| H83Q | 2 | 2 | 4 | 5 | 2 | 3 | 2 |
| Y316A | 13 | 3 | 11 | 2 | 9 | 5 | 3 |
| T338I | 7 | 5 | 3 | 42 | 3 | 4 | 4 |
| E460G | 15 | 6 | 6 | 9 | 12 | 11 | 7 |
| S783P | 12 | 18 | 24 | 8 | 8 | 15 | 10 |
| Q867L | 9 | 58 | 2 | 30 | 61 | 135 | 18 |

The initial step of the mutation selection process involved identifying the consensus leaders. The top-ranked mutation was S137P, for which all the models were highly consistent, identifying it as the most probable (Table 2). Three additional mutations were also prioritized based on the normalized average values of the substitution scores: H83Q, Y316A, and T338I. These four mutations are located in the N-terminal half of the protein. In the second step, three other mutations with high normalized average values that are distributed throughout the sequence - E460G, S783P, and Q867L-were included to explore regions associated with different sequence domains (Figure 2a).

Despite the overall similarity between models, some differences remain. For example, some mutations that appear in the top 10 after averaging are ranked outside the top 20 —or even the top 40— in individual models. The most notable example is Q867L, which is the second most probable mutation according to ESM1v_2, whereas other models rank it much lower, in some cases below position 30 or even 100. The combination of a high average score, 18 overall, and its location in a distinct sequence domain supported its inclusion as a diversity-oriented candidate in the selected set.

To further understand the potential functional impact of the selected mutations, we employed the Caver algorithm^37^ to identify the catalytic pocket of the enzyme. It predicted a main pocket including the amino acids Y96, N140, A141, E142, N199, E200, Y260, E298, and Y364. We then predicted the three-dimensional structure of the mutants using AlphaFold2 and visualized their positions relative to the catalytic pocket (Figure 2b). Notably, none of the selected mutations were located within the identified catalytic pocket. However, H83Q, S137P, Y316A, and T338I were observed to be in proximity, suggesting a potential influence on the stability of the catalytic pocket or substrate binding. Conversely, E460G, S783P, and Q867L were distant from the active site.

While mutations in active sites or substrate-binding areas may theoretically be more likely to impact catalytic efficiency, and peripheral mutations might affect stability or folding^46^, recent studies have highlighted the activity-enhancing potential of distal mutations^47–49^, potentially through conformational changes that impact activity. Understanding the specific location of each mutation provides insight into its individual contribution and highlights the potential for synergistic effects. This observation aligns with the tendency of PLMs to predict substitutions outside highly conserved active-site regions, which are crucial for maintaining enzyme function. These patterns reflect the heterogeneous biochemical and structural consequences of the substitutions (Figure S2).

### Characterization of substrate specificity and kinetic constants

Following the transformation of *K. phaffii* with the mutant β-galactosidase sequences, we successfully obtained transformants for the WT and the seven *A. oryzae* β-galactosidase variants. Initial activity screening of two clones per enzyme revealed activity in all transformed clones, while untransformed controls showed no activity (Figure S3).

Notably, the highest activity against *o*NPGal was observed in clones expressing the E460G variant, with one clone exhibiting significantly higher activity (up to 8 U/mL), likely due to gene multicopy integration^50^. The other clones displayed good activity levels in the range of 2-3 U/mL. Recombinant proteins were purified to homogeneity in a single molecular exclusion chromatography step. While the chromatogram showed up to six protein peaks (detected at A280 nm), β-galactosidase activity was exclusively detected in one fraction. SDS-PAGE analysis of purified active fractions revealed a single band for all preparations (Figure S3b), corroborating their homogeneity. The observed molecular mass of 140-150 kDa was higher than the theoretically calculated value, 108 kDa, based on the amino acid sequence, suggesting a glycosylation level of 30-40%. *K. phaffii* is known for its post-translational modification capabilities, particularly N-linked glycosylation, resulting in high-mannose-type glycosylation^51^.

Regarding the analysis of the monomeric nature of the enzymes, the major sedimentation peak observed at 6.5–6.7 S for both WT enzyme and E460G variant is consistent with the expected mass and hydrodynamic behavior of a monomeric, globular form of the protein (Figure S4). Specifically, the wild type shows a predominant peak at 6.5 S, while the mutant E460G displays a main peak at 6.7 S, further supporting the assignment to the monomer.

All tested β-galactosidases exhibited higher hydrolytic activity (*k_cat_/K_m_*) on the synthetic substrate *o*NPGal compared to the natural substrate lactose (Figure 3). This is consistent with *o*NPGal’s typically enhanced degradation due to its better leaving group^52^. The analysis of kinetic parameters allowed the assessment of the impact of individual mutations on substrate affinity and catalytic efficiency relative to the WT enzyme (Tables 3 and 4).

**Figure 3.**
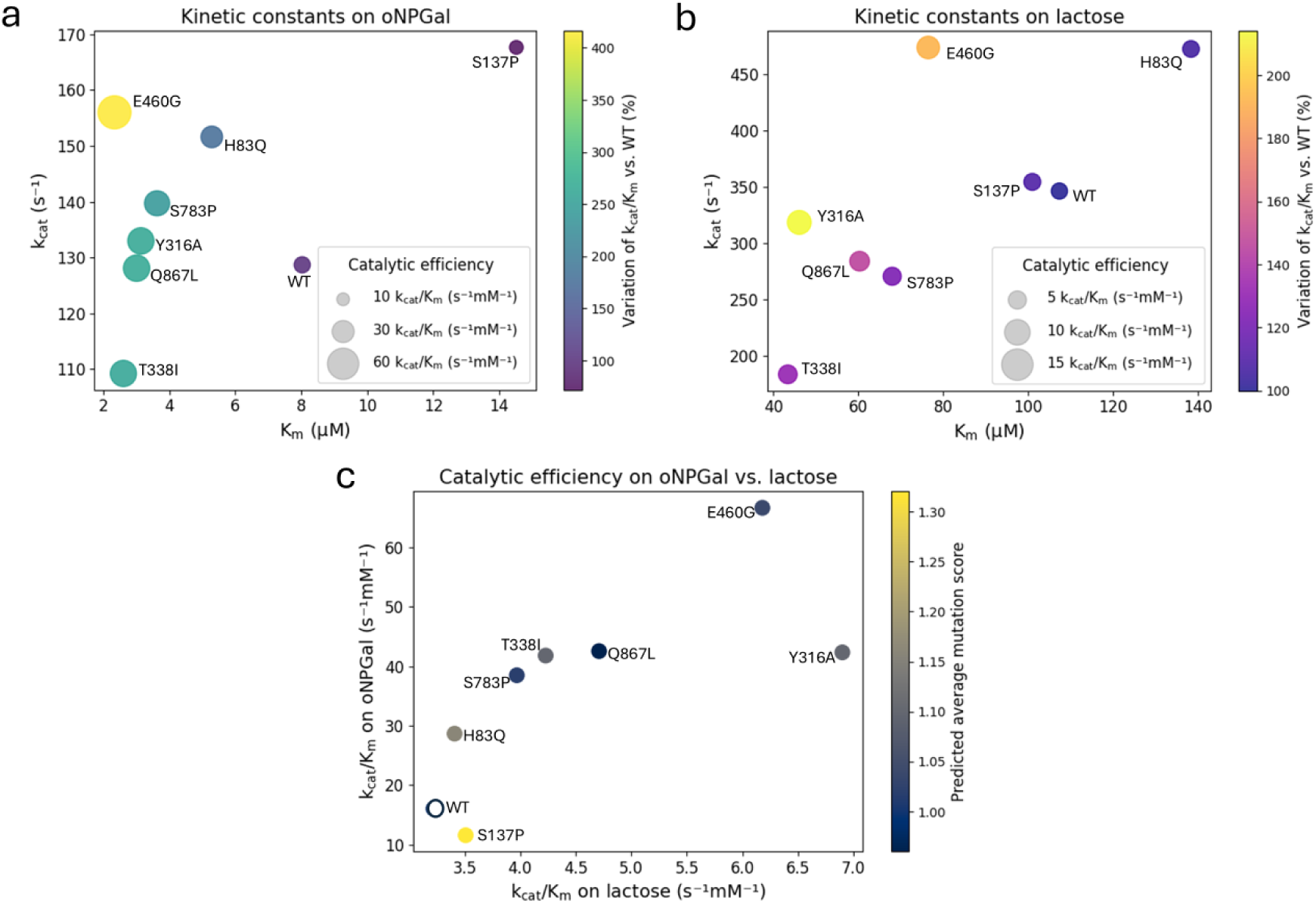
Kinetic characterization of the WT and selected mutant enzymes. (a) and (b) showcase the kinetic constants (*k_ca_*_t_ vs. *K_m_*) for the substrates *o*NPGal and lactose, respectively. For both plots, the circle size corresponds to the catalytic efficiency (*k_cat_/K_m_*) and the circle color corresponds to its variation with respect to the catalytic efficiency of the WT. (c) Comparison of catalytic efficiency (*k_cat_/K_m_*) on oNPGal vs. lactose. The color of each point represents the average normalized substitution score across the six PLMs (see Materials and Methods).

**Table 3.** Kinetic constants for the WT and 7 β-galactosidase variants acting on *o*NPGal.

| Mutation | $K_m$ (mM) | $k_{cat}$ (s<sup>-1</sup>) | $k_{cat}/K_m$ (s<sup>-1</sup>/mM) | Variation of $k_{cat}/K_m$ vs. WT (%) |
| --- | --- | --- | --- | --- |
| WT | 8.03 | 128.69 | 16.02 | 100 |
| H83Q | 5.29 | 151.61 | 28.65 | 178 |
| S137P | 14.52 | 167.66 | 11.54 | 72 |
| Y316A | 3.14 | 132.97 | 42.34 | 264 |
| T338I | 2.61 | 109.21 | 41.80 | 260 |
| E460G | 2.34 | 156.01 | 66.67 | 416 |
| S783P | 3.63 | 139.71 | 38.48 | 240 |
| Q867L | 3.01 | 128.08 | 42.55 | 265 |

**Table 4.**
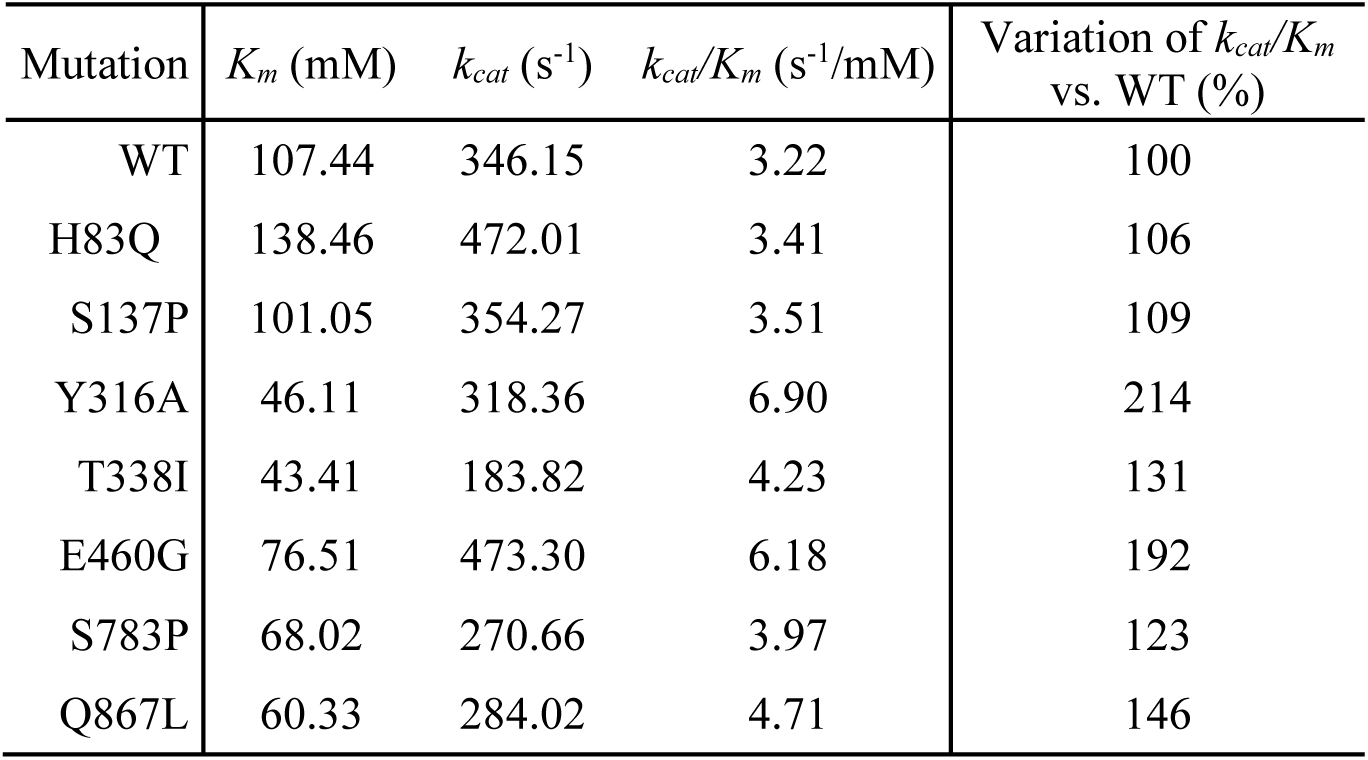
Kinetic constants for the WT and 7 β-galactosidase variants acting on lactose.

All variants, except S137P, showed improved hydrolytic activity on *o*NPGal compared to the WT enzyme. The E460G mutant displayed the most significant increase in catalytic efficiency (*k_cat_/K_m_*), achieving a four-fold improvement. Similar trends were observed for lactose degradation, with the E460G mutant showing a nearly two-fold increase in activity compared to the WT enzyme. Additionally, Y316A emerged as the most efficient variant for lactose hydrolysis, exhibiting a 2.14-fold increase in catalytic efficiency. The remaining variants also matched or surpassed the catalytic efficiency of the WT enzyme, highlighting their potential for industrial lactose degradation.

Figures 3c and S5 show the relationship between hydrolytic activity across the two different substrates, as well as the average mutation score for each variant. As previously described, there is no direct correlation between high score and high activity, since PLMs were not specifically trained to optimize any particular property. Nevertheless, all the mutations selected based on PLM scores resulted in stable enzymes that improved the hydrolytic activity of the original enzyme, something that would unlikely have occurred with random mutagenesis^53^.

The most substantial improvement in catalytic efficiency was conferred by the distal E460G mutation. Distal mutations, located far from the active site, can enhance enzyme activity by stabilizing the overall protein structure, improving substrate binding, or facilitating optimal conformational changes during catalysis^54,55^. However, the second most effective mutation, Y316A, was located near the catalytic pocket, demonstrating the pipeline’s ability to identify beneficial variations even in evolutionary conserved regions.

To better understand the molecular drivers behind these efficiency gains, we analyzed the specific physicochemical shifts introduced by the top-performing mutations (Figure S2). For Y316A, situated in the immediate vicinity of the catalytic pocket (Figure 2b), the substitution replaces a bulky, aromatic tyrosine (163Å^3^) with a smaller, hydrophobic alanine (67Å^3^). This drastic reduction in steric volume could alleviate spatial constraints near the active site, potentially streamlining substrate access or product release. In contrast, the distal E460G mutation replaces a negatively charged glutamic acid with glycine. This modification eliminates a local charge interaction and, given glycine’s high conformational freedom, likely modulates local backbone flexibility, a factor often linked to enhanced catalytic dynamics in distal regions.

### Physicochemical properties of the enzymes

All enzyme variants displayed similar optimal pH values, with maximal activity around pH 5-6 (Figures S9 and S10). No major shifts in optimum pH were detected among the mutants compared to the WT, indicating that the substitutions predicted by the PLM models did not alter the protonation requirements of catalytic or substrate binding residues. In addition, pH stability profiles were broadly conserved across the panel: all enzymes retained close to 100% activity after 24 h between pH 3 and 7, and the most stable variants showed slightly improved tolerance at pH 2 and pH 8 (data not shown), although no large differences were observed overall. These results suggest that the engineered mutations preserve the native acidic behavior of the *A. oryzae* β-galactosidase and maintain functional activity across a wide pH range, which is relevant for food-processing applications.

Regarding thermal stability, after 24h of incubation all variants tolerated thermal stress at 50 °C and exhibited gradual loss of activity above that temperature, consistent with the known moderate thermal tolerance of fungal β-galactosidases. However, one mutation, T338I, stood out by maintaining approximately 80% of its initial activity after 24 h at temperatures up to 60 °C. The S137P and E460G variants also displayed slightly improved stability relative to the wild-type, but their effect was clearly less pronounced than that of T338I (Figures S11 and S12). This pattern is particularly interesting in light of the structural context of the mutations: T338 is located close to the catalytic region, and its replacement by a bulkier, more hydrophobic isoleucine likely strengthens local packing interactions in this area, providing additional resistance to thermal denaturation. In contrast, E460G is distal from the active site and may exert subtler long-range effects on protein dynamics, consistent with its more moderate stabilizing impact.

Taken together, these data show that the PLM-derived substitutions not only improve catalytic efficiency but can also modulate the overall stability of the enzyme, with T338I standing out as a mutation that enhances long-term thermostability without compromising activity in the acidic pH range. This is a particularly valuable trait for industrial lactose hydrolysis, where prolonged operation at elevated temperatures is often desirable to increase substrate solubility, reduce contamination risk and improve process robustness. The fact that such a stabilizing mutation was identified within a small set of seven experimentally tested variants reinforces the notion that PLM-guided design can efficiently enrich substitutions that simultaneously support activity and stability, thereby reducing the number of wet-lab candidates that need to be screened.

### Analysis of GOS production

A notable challenge for industrial applications of β-galactosidases is their potential to exhibit synthesis activity, producing co-products like galactooligosaccharides and other oligosaccharides^29^. However, GOS production is valuable due to their recognized prebiotic properties^31,32^. The *A. oryzae* β-galactosidase is well-documented for its significant transgalactosylation activity, prompting us to determine whether any of the seven variants can also improve its synthesis capacity. Prior protein engineering has successfully enhanced GOS production yields in other β-galactosidases^40^.

To assess the enzyme variants’ ability to catalyze galactose transfer to various acceptor molecules and form galactooligosaccharides, we performed transgalactosylation reactions under controlled conditions. Based on elution behavior and previously published GOS-profiles for β-galactosidases ^56–58^, we tentatively assigned peaks 1 and 2 in the transglycosylation chromatograms to galactooligosaccharides. The resulting GOS products were quantified using high-performance liquid chromatography (HPLC) (Figure S6a). The transgalactosylation activity showed minimal differences among the tested enzyme variants and relative to the WT enzyme (Figure 4). These results indicate that the evaluated mutations generally do not significantly affect GOS synthesis. However, the E460G substitution appears to be an exception, showing a potential negative correlation between the improved lactose degradation activity and a reduction in GOS synthesis (Figure S6b). This suggests that residue E460 might play a role in modulating the balance between these two activities. The transgalactosylation potential of fungal β-galactosidases, as observed in this study, raises important questions regarding the physiological relevance and evolutionary trajectory of this catalytic trait. β-galactosidases from *A. oryzae* are classically described as hydrolytic enzymes, primarily tasked with the breakdown of galactosyl-containing oligosaccharides and polysaccharides derived from plant cell walls, such as galactomannans and arabinogalactans^59^. This hydrolytic activity supports fungal saprotrophy. In contrast, transgalactosylation has little, if any, documented physiological function in natural ecosystems. Rather, it is widely regarded as a catalytically incidental activity^60^. This side activity becomes particularly evident under high substrate concentrations and low water activity; conditions not typically encountered in natural niches but frequently replicated in industrial processes designed to favor synthetic over hydrolytic processes^61^.

**Figure 4.**
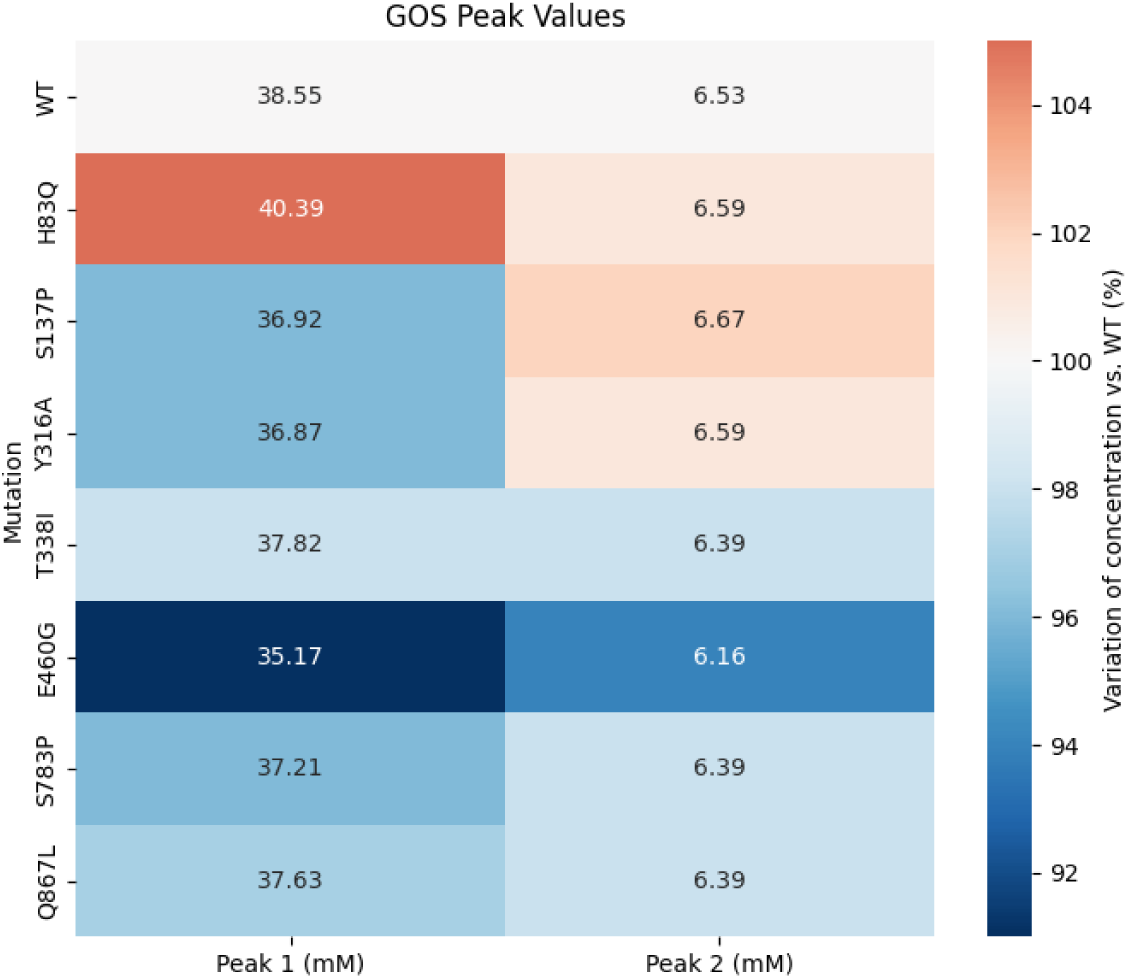
Maximum concentrations of synthesis products obtained in the transgalactosylation reactions catalyzed by the β-galactosidases. The concentrations (mM) refer to lactose equivalents for each product. The color indicates the variation relative to the WT.

In this context, the results of our study highlight how structural determinants can influence the balance between competing reaction mechanisms. The reduction in GOS yield associated with the E460G mutation, despite improved lactose hydrolysis, suggests that this residue may contribute to modulating water versus acceptor sugar access. The increased flexibility introduced by the glycine substitution (E460G) might favor the entry of the smaller water molecule (hydrolysis) over the bulkier lactose acceptor (transgalactosylation). From an evolutionary perspective, the presence of latent transgalactosylation activity in *A. oryzae* β-galactosidases can be interpreted as a case of catalytic promiscuity^62^. While PLMs effectively optimized the ‘fitness’ of the enzyme, they do not inherently distinguish between reaction pathways. Notably, nature has evolved dedicated galactosyltransferases to fulfill anabolic roles in glycan biosynthesis, indicating that synthetic functions are not incompatible with galactosyl donor chemistry, but may require more specialized active site architectures than those found in canonical β-galactosidases^63^. Further structural and kinetic studies with β-galactosidase will be necessary to fully elucidate the molecular basis of this trade-off, increase GOS production, and to harness it for rational enzyme design.

Finally, it is worth noting that the mutation most consistently predicted as highly probable across all models, S137P, did not emerge as one of the top variants in terms of GOS formation or lactose hydrolysis in our assays. Rather than indicating a limitation, this highlights the fact that the used PLMs capture general evolutionary and structural preferences but are not explicitly optimized for every single enzymatic property. Consequently, a mutation such as S137P may be highly compatible with the native structural and evolutionary constraints of the protein while influencing properties different from those measured here. For example, such substitutions may enhance attributes like conformational rigidity, or expression efficiency.

## CONCLUSIONS

Protein Language Models, despite their recent introduction to the protein field, have already demonstrated significant impact on enzyme optimization. Whereas supervised PLM pipelines typically require rounds of experimental characterization to fit a regressor linking embeddings to activity, our results show that the ensemble-based zero-shot strategy implemented here — averaging the normalized score distributions across six PLMs — can identify beneficial substitutions in an industrially relevant enzyme with a single round of experimental validation. All seven predicted substitutions yielded stable, expressible variants, and all but one matched or surpassed the wild-type’s catalytic efficiency on lactose. This level of predictive enrichment is substantially higher than what is typically achieved through random mutagenesis and follows from three design choices of our pipeline: (i) the formal conciliation of six independently trained PLMs, which mitigates individual model biases; (ii) a two-step, diversity-aware selection that combines top consensus leaders with high-scoring candidates from under-represented sequence domains, rather than relying on a purely top-ranked consensus; and (iii) the demonstration that a single round of zero-shot inference can simultaneously surface variants with improved catalytic efficiency and variants with markedly enhanced thermostability in an industrially deployed enzyme. This success could be due to specific circumstances related to the β-galactosidase case study, but it also suggests that the use of several PLM and the use of a conciliation criterion implemented in our pipeline may be a valuable strategy for optimizing enzymes from different families, reducing the need for extensive experimental validation of predicted variants.

Additionally, this study identified novel and relevant mutations for β-galactosidase improvement, including Y316A and E460G, which significantly increased the enzyme’s hydrolytic activity by up to 4.16-fold individually, underscoring the effectiveness of our optimization process. Interestingly, this enhanced hydrolytic activity was not mirrored in the transgalactosylation activity, raising questions about the evolutionary relationship between these two enzymatic functions.

This work highlights the potential of AI-assisted enzyme evolution platforms to not only design more efficient enzyme variants but also to unlock new possibilities for industrial applications. The strategic application of PLMs has the potential to transform industrial biocatalysis by offering more efficient, cost-effective, and tailored enzymatic solutions to meet the evolving demands of modern industry.

## Supporting Information

Sequences used in this study (Table S1), check of linearity of the reactions at three different times (Table S2), consensus score for each candidate substitution across the ensemble (Table S3), mutation score heatmaps, residue substitution overview, activity screens, SDS-PAGE gels, ultracentrifugation data, and HPLC chromatograms, Michaelis-Menten plots of the kinetics, and pH and temperature assays (Figures S1-S12)

## Conflict of interests

A patent application covering aspects of the computational pipeline and derived enzyme variants described in this manuscript has been filed. This intellectual property context determines the terms of code availability described in the Code and Data Availability section.

## Acknowledgements

The authors would like to thank the members of Barriuso’s laboratory and TheNextPangea for their insightful discussions and technical assistance in this paper. We would also like to thank the Molecular Interactions Facility from CIB-CSIC for performing the analytical ultracentrifugation experiments. The authors acknowledge the support toward the publication fee by the CSIC Open Access Publication Support Initiative through its Unit of Information Resources for Research (URICI). This work was supported by the Principality of Asturias (SEKUENS PID reference IDE/2023/000449).

## Code and Data Availability

The computational pipeline developed in this study — comprising per-position substitution score extraction from ESM2 and ESM1v, conciliation across the six-model ensemble, and candidate ranking with diversity-aware selection — is available from the corresponding authors upon reasonable request, subject to the intellectual property considerations described in the Conflict of Interests section. The ESM2 and ESM1v model checkpoints used in this work are publicly available through their original publications ^35,36^. Consensus substitution scores for each candidate mutation in the *A. oryzae* β-galactosidase wild-type sequence (UniProt Q2UCU3) are provided as Supporting Information (Table S3). These data support Tables 1 and 2 and the selection of the seven experimentally validated variants. The nucleotide sequences of all mutant constructs generated in this study are included in Table S1.

## Author contributions

A.T.F. and A.F. designed the computational framework, performed PLM-based predictions and the *in-silico* validation. J.A.M-L. and A.P. designed the enzyme validation workflow. J.A.M-L. performed enzyme production and activity evaluation. A.T.F., A.F., J.A.M-L and A.P. participated in results’ discussion and manuscript writing. J.B. and F.G.O. designed the study and wrote the manuscript.

