## Supplementary Figures for "AI-assisted improvement of *Aspergillus oryzae* β-galactosidase using an Ensemble of Protein Language Models"

**Figure S1**. Heatmap of average predicted mutation scores across the protein sequence. Each cell represents the predicted average score of a single amino acid substitution at a given sequence position.

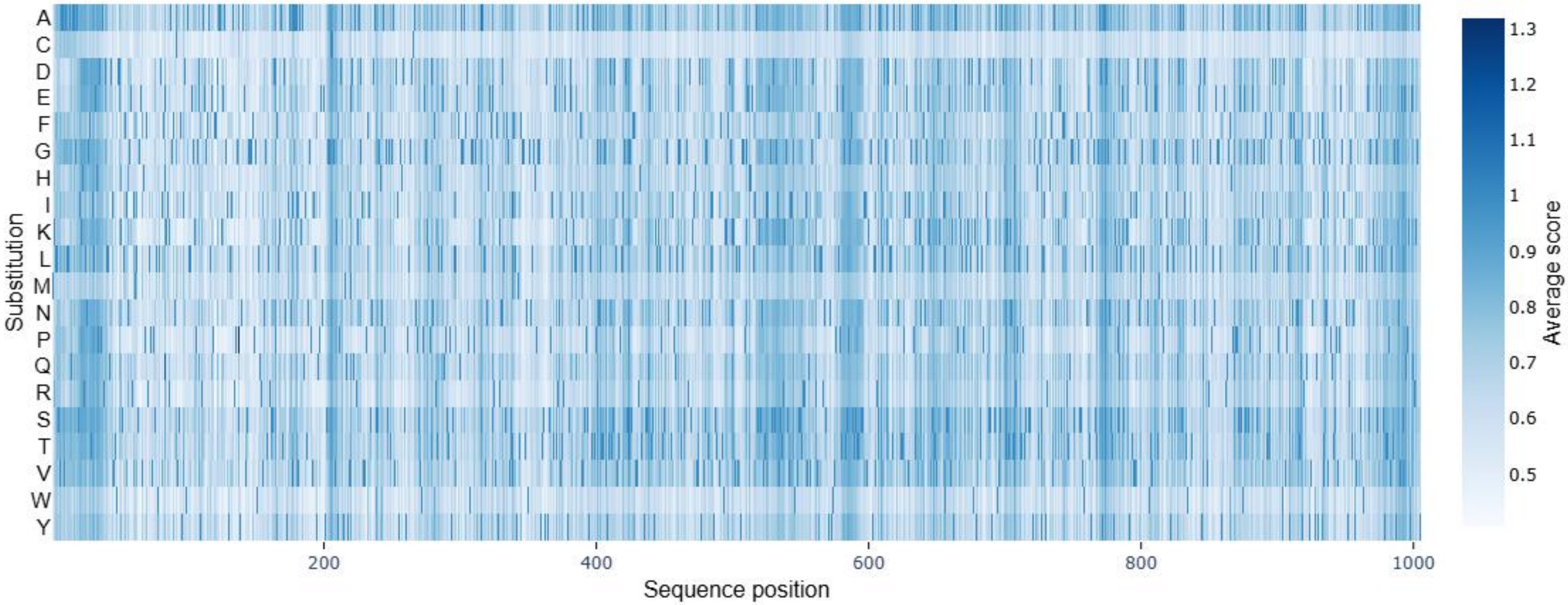

| | WT Residue | | | | Mutant | | | |
|---|---|---|---|---|---|---|---|---|
| **H83Q** | **Histidine** | Positively charged or neutral | 118 Å$^3$ | 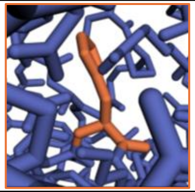 | **Glutamine** | Uncharged | 143 Å$^3$ | 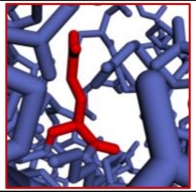 |
| **S137P** | **Serine** | Uncharged | 73 Å$^3$ | 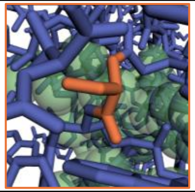 | **Proline** | Uncharged | 90 Å$^3$ | 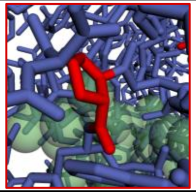 |
| **Y316A** | **Tyrosine** | Uncharged | 163 Å$^3$ | 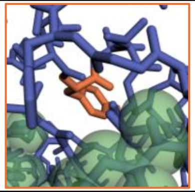 | **Alanine** | Uncharged | 67 Å$^3$ | 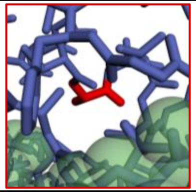 |
| **T338I** | **Threonine** | Uncharged | 93 Å$^3$ | 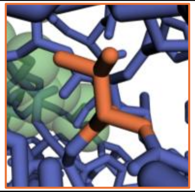 | **Isoleucine** | Uncharged | 124 Å$^3$ | 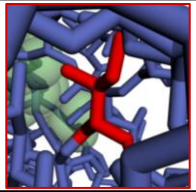 |
| **E460G** | **Glutamic Acid** | Negatively charged | 138 Å$^3$ | 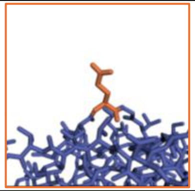 | **Glycine** | Uncharged | 48 Å$^3$ | 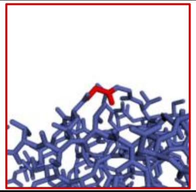 |
| **S783P** | **Serine** | Uncharged | 73 Å$^3$ | 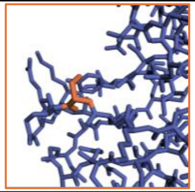 | **Proline** | Uncharged | 90 Å$^3$ | 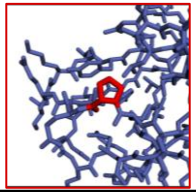 |
| **Q867L** | **Glutamine** | Uncharged | 143 Å$^3$ | 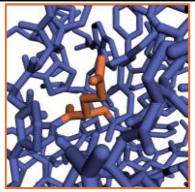 | **Leucine** | Uncharged | 124 Å$^3$ | 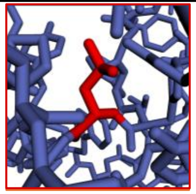 |

**Figure S2**. Comparison of wild-type (WT) and mutant residues showing amino acid identity, charge state, approximate side-chain volume (Å³), structural snapshot and physicochemical classification.

- Nonpolar hydrophobic
- Polar hydrophilic, uncharged
- Nonpolar hydrophobic, Aromatic
- Polar hydrophilic, charged

a

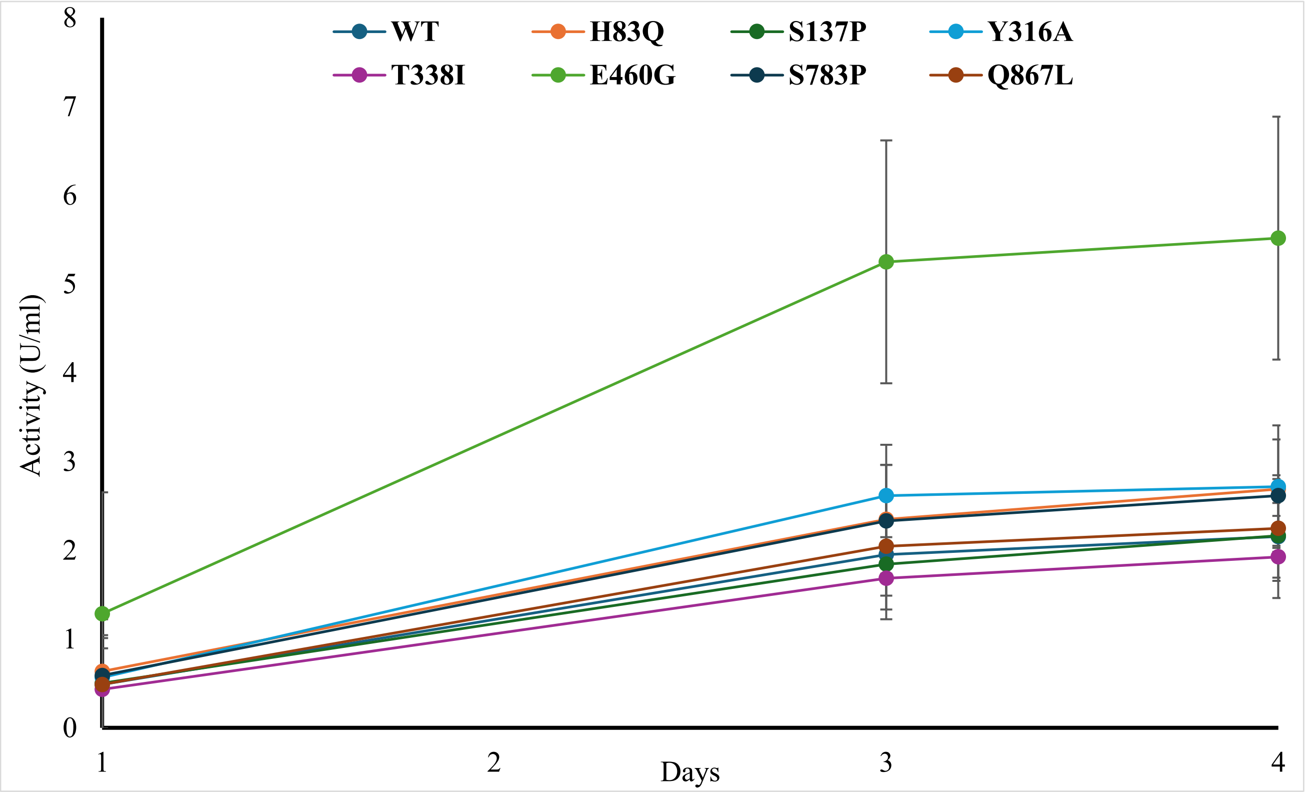


b

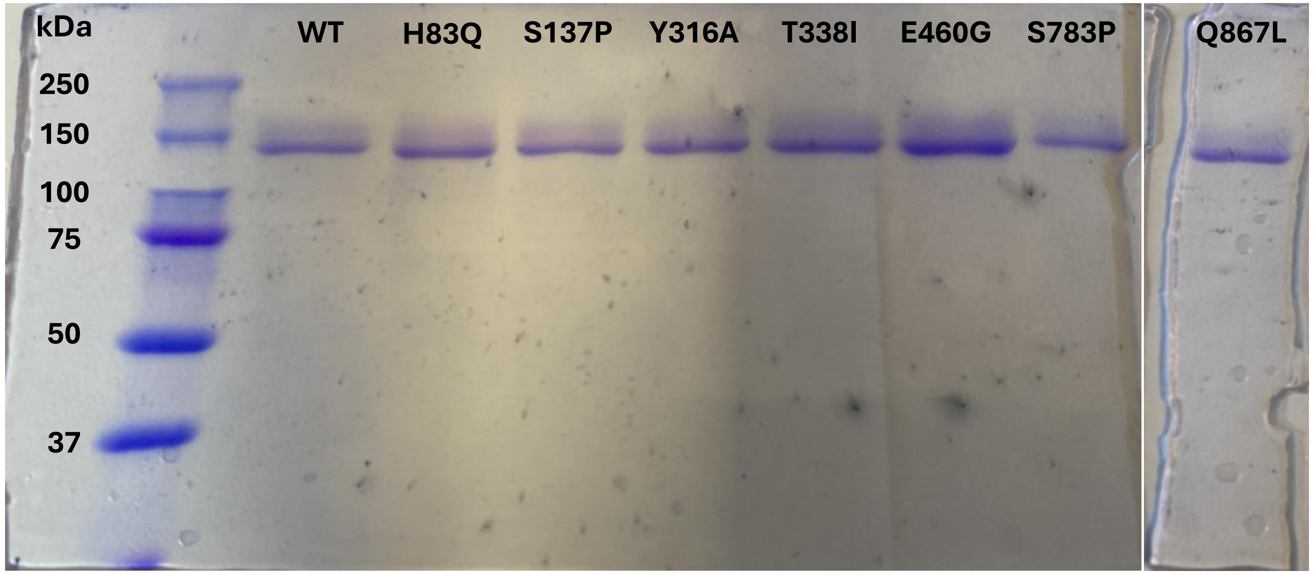


**Figure S3**. (a) β-Galactosidase activity of secreted β-galactosidase, measured using *o*NPGal as substrate, of the *K. phaffii* strains generated in this study. (b) SDS-PAGE analysis of the β-galactosidases used in this study.

**Figure S4.** Sedimentation velocity assay of the E460G variant into analytical ultracentrifugation cells. The monomeric form of the enzyme is detected at 6.7 S corresponding to 114 kDa.

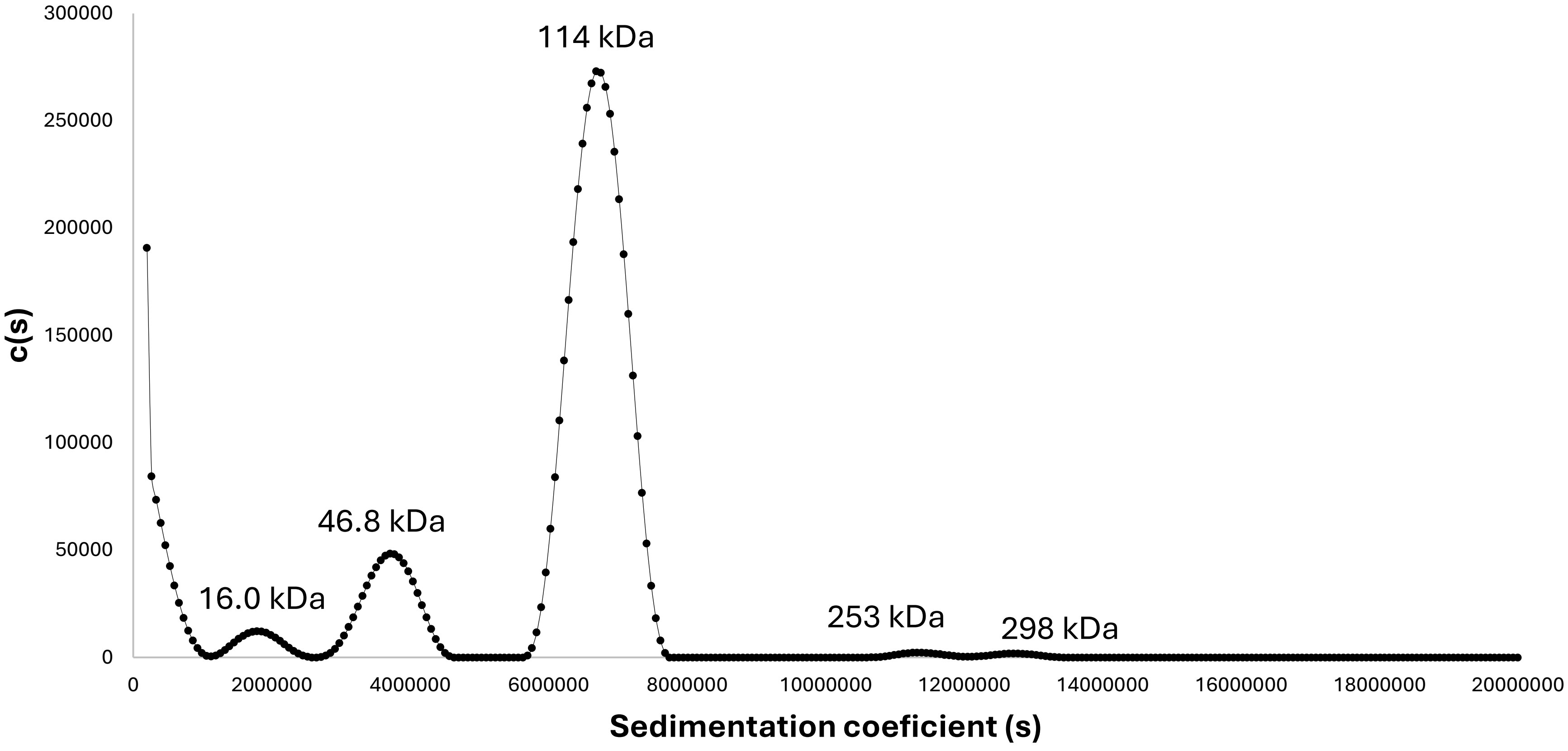

**Figure S5.** Relationship between predicted average mutation score and catalytic efficiency ($k_{cat}/K_m$) for mutant variants. (a) Catalytic efficiency on *o*NPGal. (b) Catalytic efficiency on lactose.

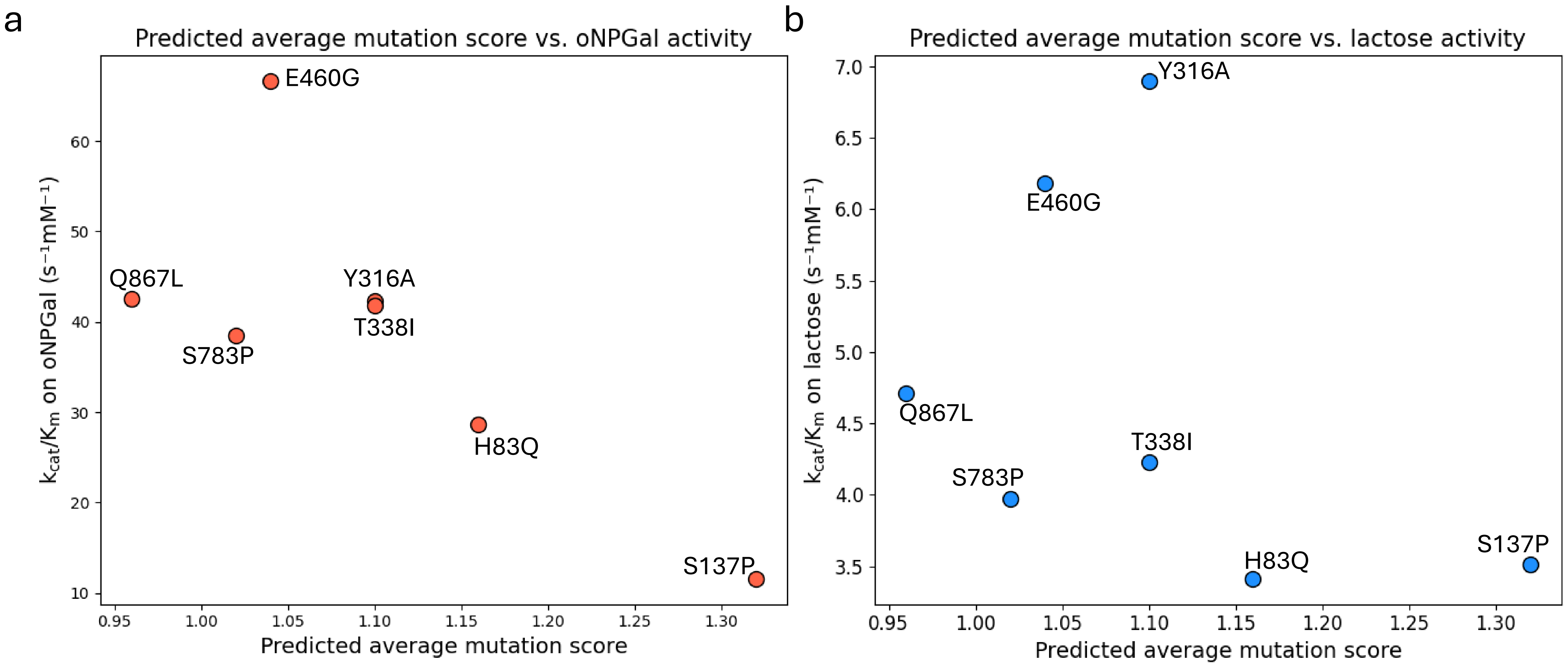

**Figure S6.** (a) Chromatographic profile of the analysis of transgalactosylation reactions. As an example, the profile observed after 2 h of reaction with the H83Q enzyme is shown. (b) Comparison of GOS production and catalytic hydrolysis efficiency ($k_{cat}/K_m$) for the WT and the seven selected variants. Staked bars represent GOS production, with Peak 1 and Peak 2 in light blue and dark blue, respectively. The red line indicates the catalytic efficiency ($k_{cat}/K_m$) of each variant.

a b

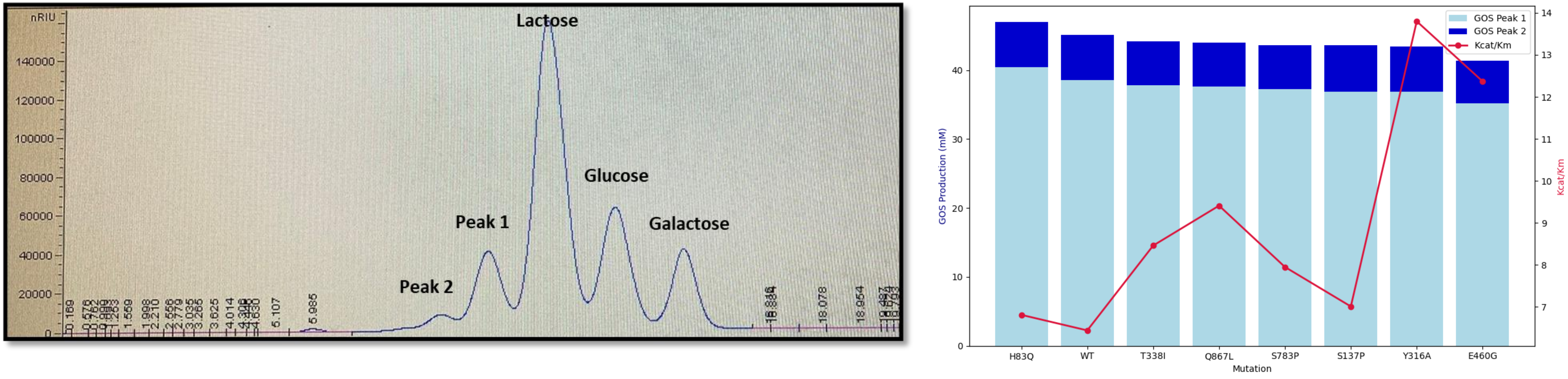

**Figure S7**. Michaelis-Menten plots of reactions over *o*NPGal

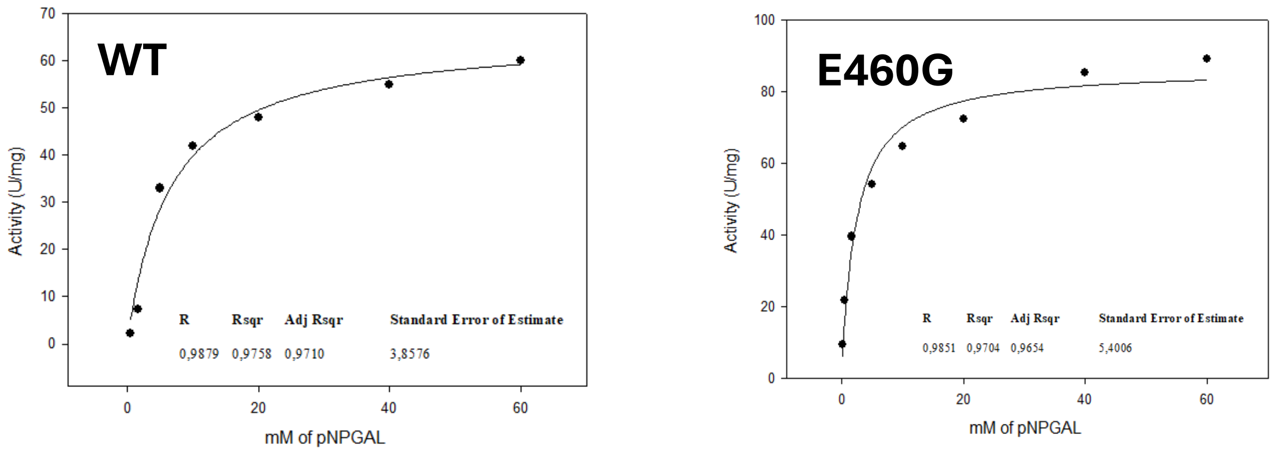


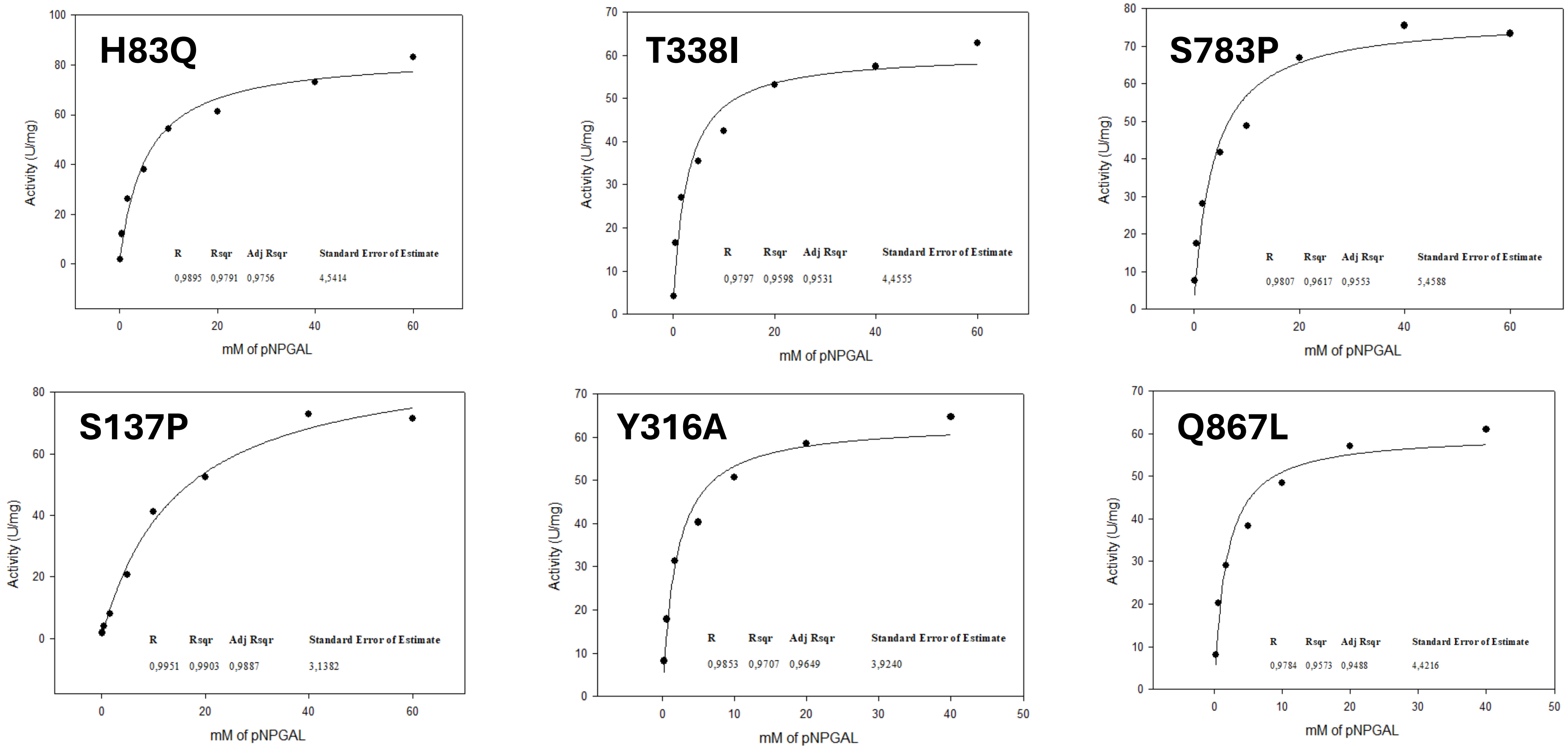

**Figure S8**. Michaelis-Menten plots of reactions over lactose

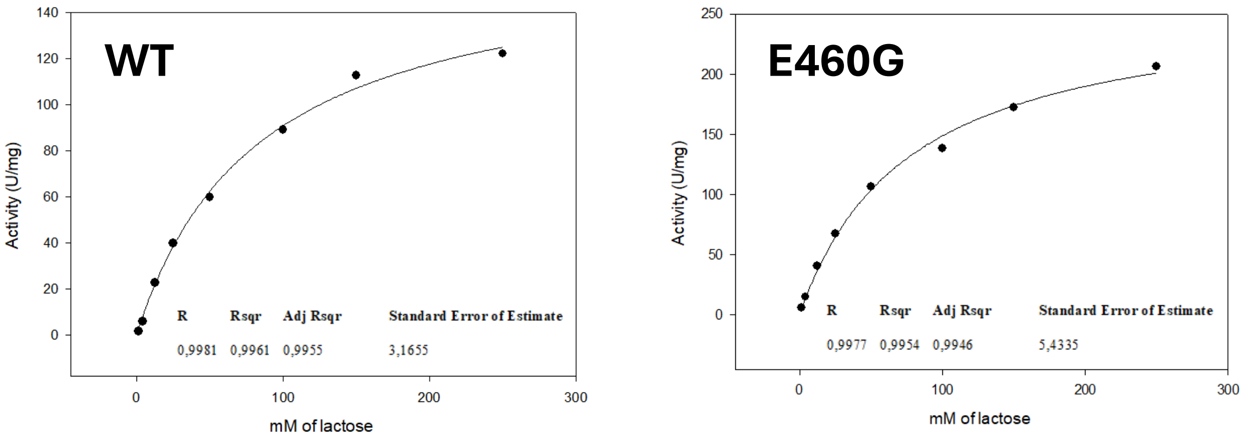


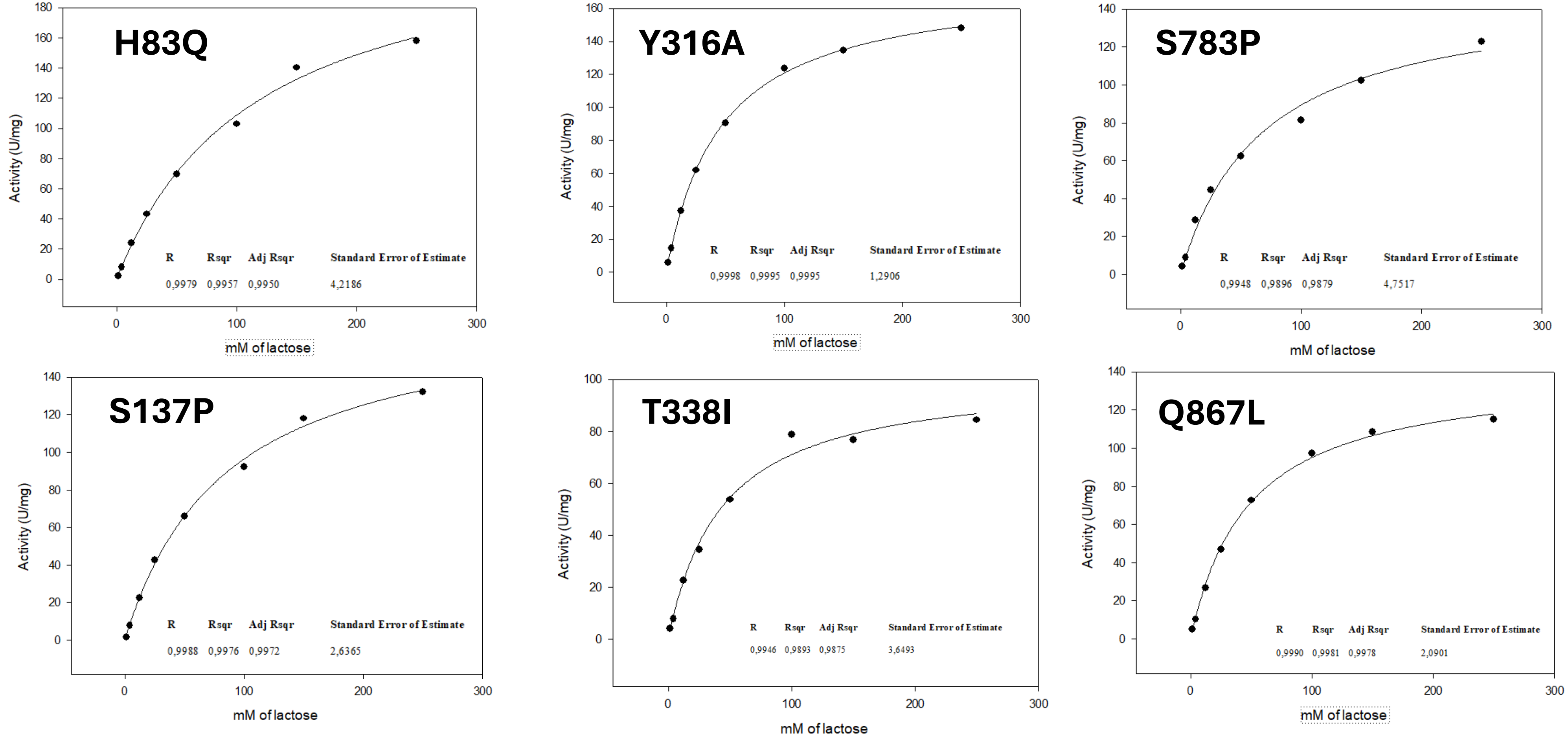

**Figure S9.** Optimal pH of WT enzyme and H83Q, S137P and Y316A variants.

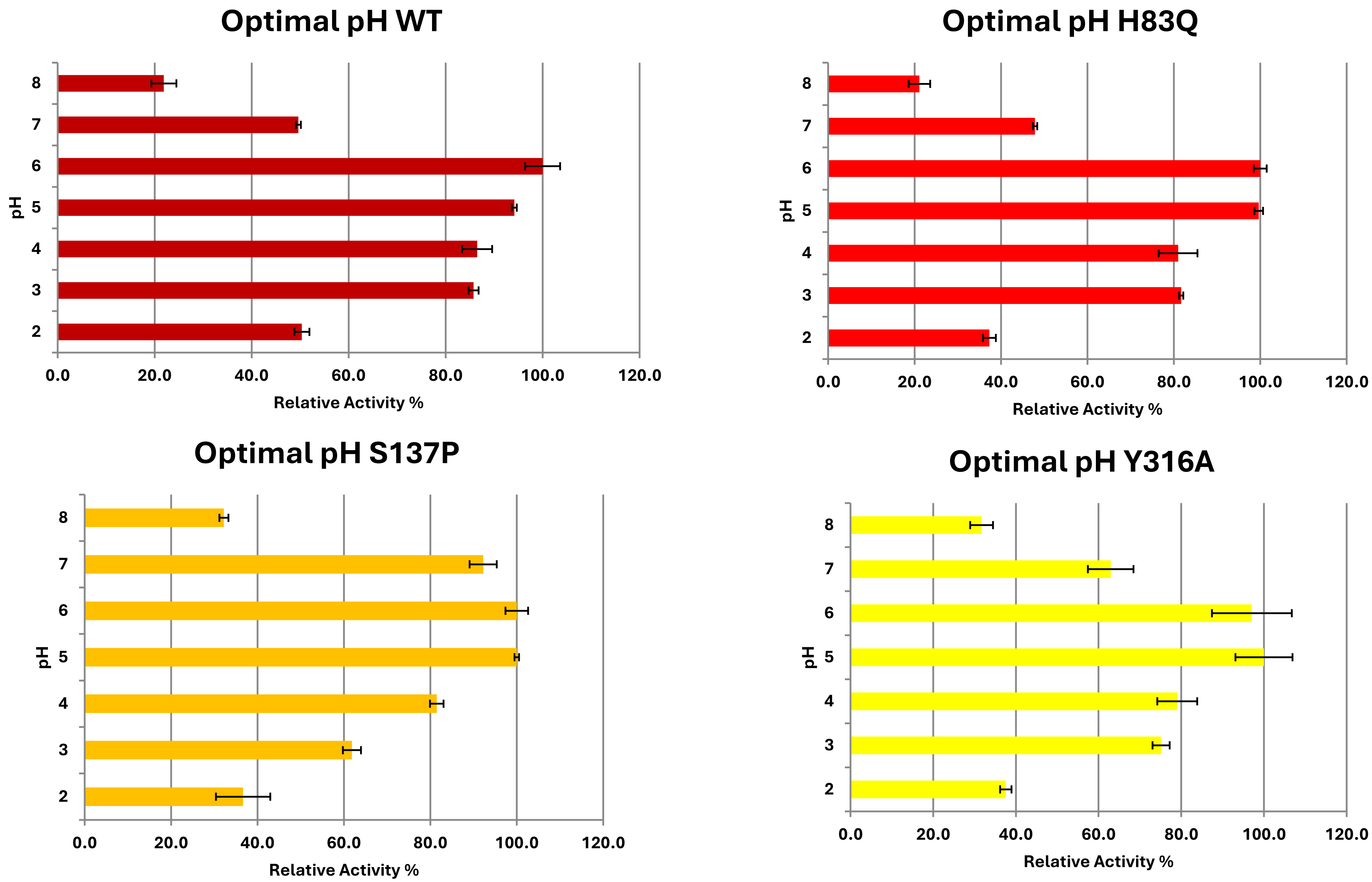

**Figure S10.** Optimal pH of T338I, E460G, S783P and Q867L variants.

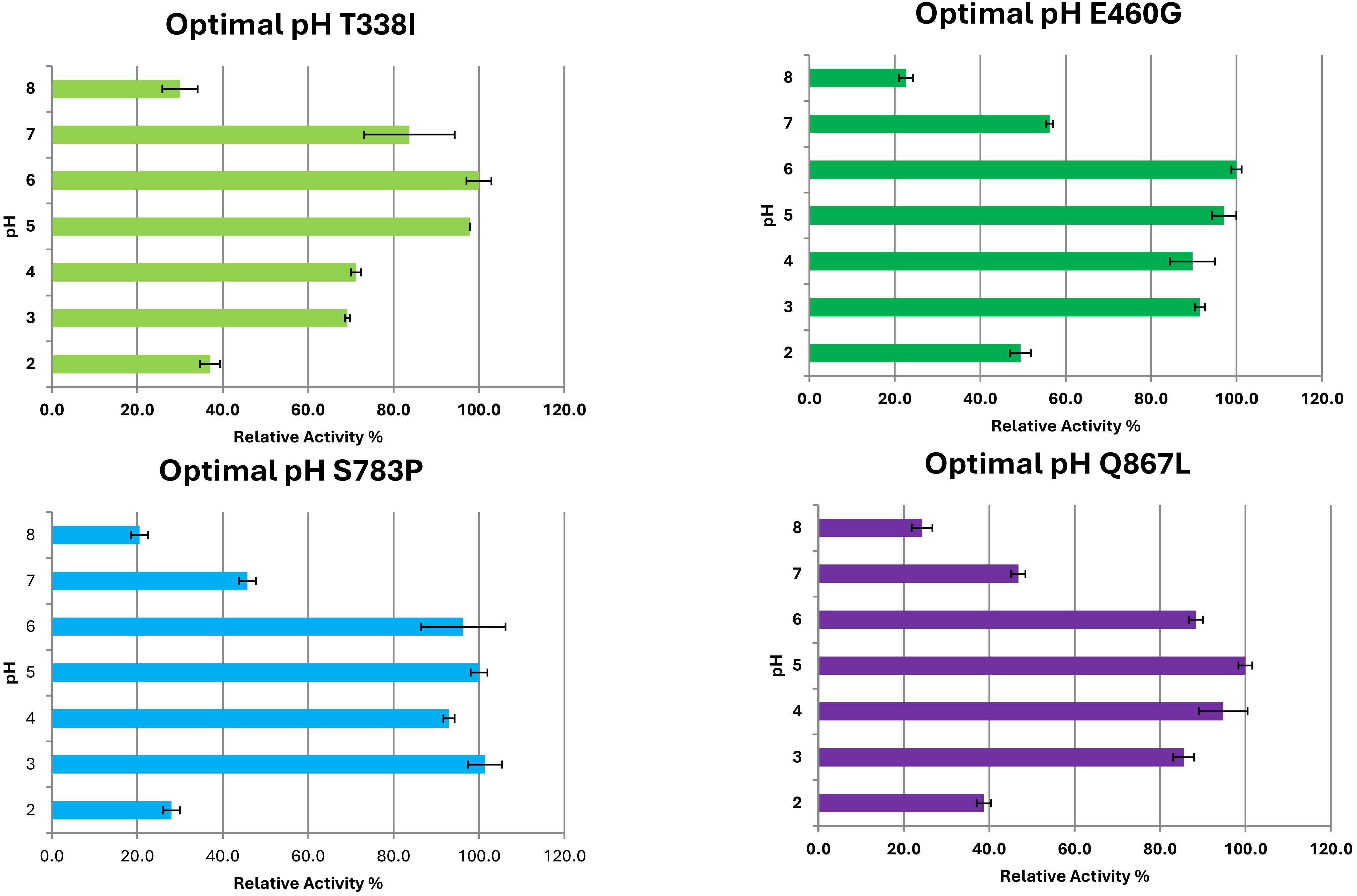

**Figure S11.** Temperature stability assay of WT enzyme and H83Q, S137P and Y316A variants.

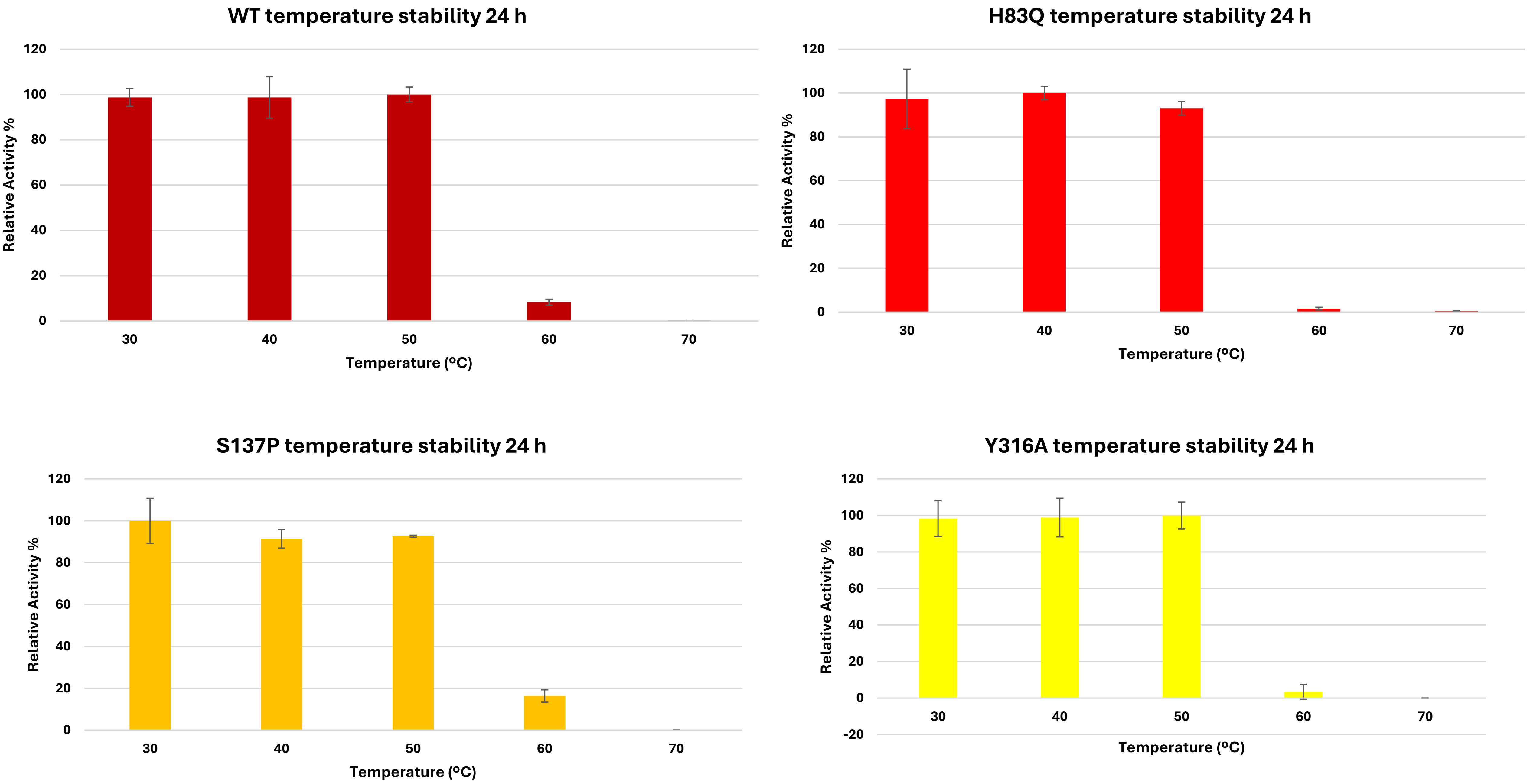

**Figure S12.** Temperature stability assay of T338I, E460G, S783P and Q867L variants.

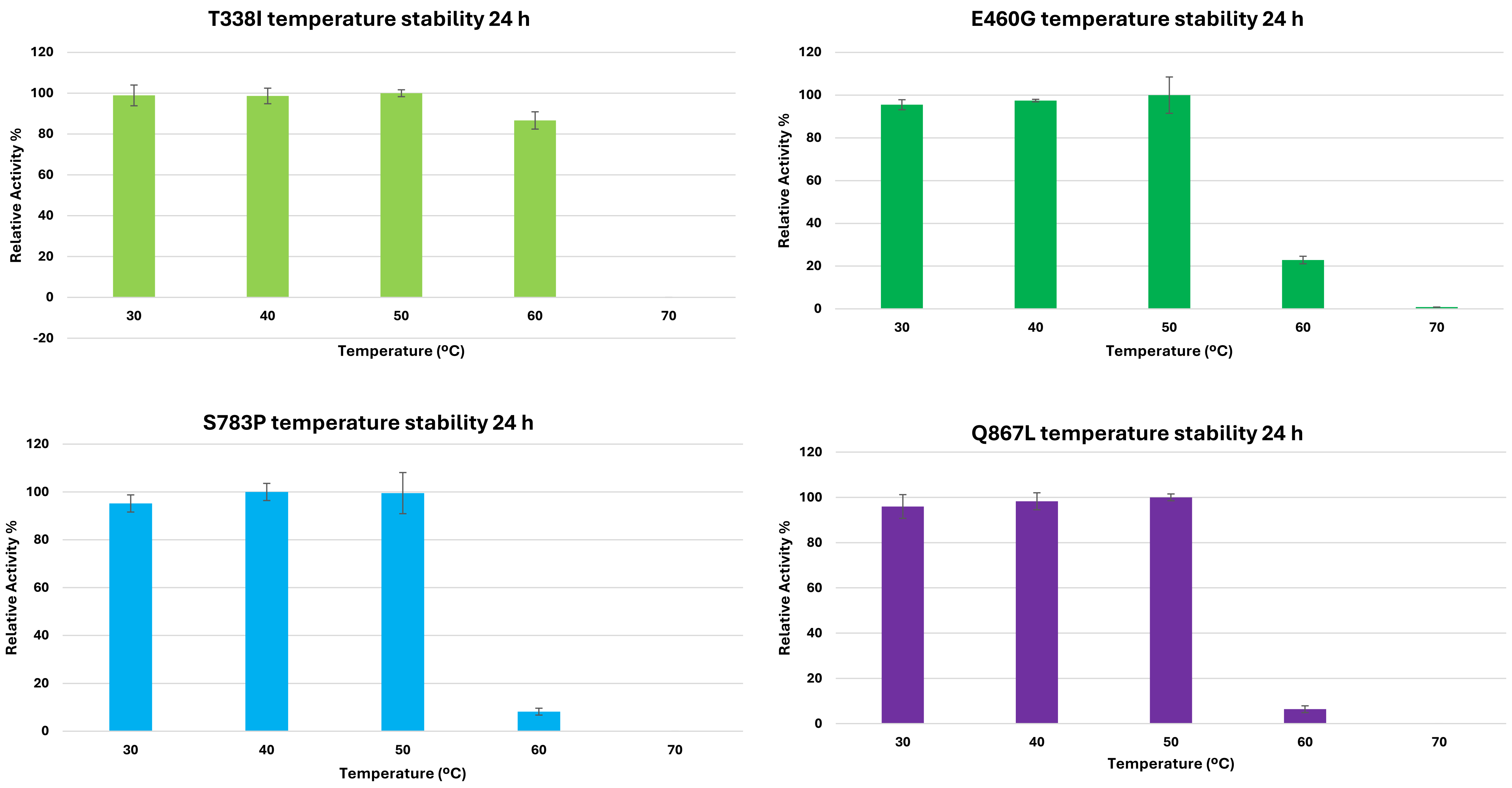
